# Identification and characterization of anti-epileptic compounds from Papaver somniferum using quantification techniques (GC-MS, FTIR), integrated network pharmacology, molecular docking, and molecular dynamics simulations

**DOI:** 10.1101/2024.07.29.605558

**Authors:** Kiran Shahzadi, Danish Rasool, Faiza Irshad, Ammara khalid, Sehar Aslam, Aisha Nazir

**Affiliations:** Departement of Bioinformatics and Biotechnology, Government College University Faisalabad; Department of Bioinformatics and Biosciences Capital University of Science and Technology Islamabad; Environmental Biotechnology Laboratory (F4) Institute of Botany, University of Punjab Lahore; Department of Biotechnology University of agriculture Faisalabad

**Keywords:** Papaver somniferum, network pharmacology, molecular dynamics, epilepsy, target prediction, molecular docking, systems pharmacology

## Abstract

Epilepsy is a common neurological condition identified by repetitive seizures that affect the overall quality of life. Existing anti-epileptic drugs have undesirable side effects, necessitating safer alternatives. This study develops an integrated computational framework to discover potential anti-epileptic leads from *Papaver somniferum* (opium poppy). Literature and databases were mined to compile all chemicals from Papaver somniferum. PubChem provided structural data, and compounds satisfying drug-likeness and bioavailability criteria were selected. GeneCards, DisGeNET and SwissTargetPrediction identified 344 target genes of the compounds and common targets with epilepsy. Network pharmacology analyses were performed. Cytoscape constructed a compound-target network comprising 5 active constituents and 22 shared targets. Degree distributions revealed molecular interactions. STRING elucidated target connectivity. Hub targets were identified using CytoHubba. GO and KEGG enrichment on 123 targets recognized biological roles and pathways. DAVID and Hiplot characterized functional annotations. Cytoscape visualized a compound-target-pathway association network involving targets, pathways, and compounds related to epilepsy. GC-MS identified 25 compounds in the Papaver somniferum extract. FTIR characterized functional groups. Molecular docking scored compound affinities for 10 targets. Autodock Vina docked 15 constituents into binding pockets. Interactions were validated using Desmond MD simulations of IL6 with scoulerine over 100 ns, assessing RMSD, RMSF, interactions. RMSD/RMSF plots and histograms characterized protein/ligand stability and flexibility. This integrative *Insilico* and *Invitro* framework facilitates prioritizing Papaver somniferum constituents for epilepsy. Network analyses provided systems-level understanding of multi-target mechanisms. Molecular modeling established structure-activity relationships, validating predicted interactions. Compounds with good ADMET profiles, network centrality, docking scores, and stable simulations emerge as candidates warranting further examination for safer anti-epileptic therapy.

## Introduction

Medicinal herbs have been used for a variety of reasons since the dawn of human history (Alves et al., 2023). A 5000-year-old Sumerian clay slab from Nagpur contains the oldest documented written account of using plants for medical purposes (Alyami et al., 2022). Because they generate a wide variety of metabolites and because there are over 250,000 different kinds of plants, they are important (Andres-Mach et al., 2021). They have the same chemical makeup—primary metabolites—that is necessary for a cell to remain alive (Araujo et al., 2022). Furthermore, a diverse array of phytochemicals and secondary metabolites found in them are implicated in interactions among different organisms (Atmaca et al., 2019). A natural chemical is created by the primary and/or secondary metabolic processes of living things (Baeeri et al., 2020). These chemicals typically have pharmacological qualities that make them useful in the treatment of a variety of illnesses. They are frequently employed as beginning points in the development of new drugs, from which analogs with superior efficacy, safety, and purity are derived (Chantarat et al., 2024). Medicinal plants produce large amounts of secondary metabolites, including flavonoids, alkaloids, phenolics, and tannins. These substances have the capacity to strengthen the innate immune system, promote resistance to disease, and promote growth in humans (Correa Basurto et al., 2023). Within the Papaveraceae family, Papaver somniferum L., also known as opium poppy, is a major medicinal plant that is best known for making opium gum (DeFreitas et al., 2022), #0}. Five significant alkaloids are included in the capsular latex of *Papaver somniferum L*. and are known as opium gum: morphine, codeine, thebaine, papaverine, and noscapine. The key substrate for the synthesis of semi-synthetic opioid alkaloids and natural opioids (such morphine and codeine) is thebaine, one of these alkaloids. Painkillers (analgesics), cough suppressants, and medications that relax smooth muscles are only a few of the medicinal uses for opium alkaloids (Dervis et al., 2024). Particularly, morphine is frequently used to treat severe pain. Epilepsy disease (PD) is a neurological condition that worsens over time and is primarily characterized by motor impairments, such as tremors, rigidity, bradykinesia, and trouble maintaining balance and posture (Domingos et al., 2024; Jiao et al., 2022).

The brain’s neural synchronization and hyperexcitability are the main causes of epilepsy symptoms (Dervis et al., 2024; Domingos et al., 2024; Jiashuo et al., 2022). This results in seizures, which can range from extended bouts of intense shaking to shorter, almost imperceptible episodes. A common neurological disorder that affects people of all ages is epilepsy. It is typified by repeated seizures, which are brief bursts of involuntary movement that can affect the whole body (generalized) or just a portion of it. They are frequently followed by a transient impairment in mental function. Epilepsy is one of the most prevalent neurological disorders in the world, affecting about 50 million individuals (Domingos et al., 2024; L. Z. Li et al., 2023).

Anti-seizure drugs like levetiracetam, valproate, lamotrigine, etc. are currently used as treatments for epilepsy. However, these medications’ adverse effects can result in weariness, sleepiness, nausea, and behavioral abnormalities (Engel & Pitkanen, 2020; S. Li et al., 2023). This accentuates the necessity of exploring new therapeutic avenues for the management of epilepsy. Consequently, herbal medicine is a successful and affordable anti-epileptic therapy choice. In 2007, network pharmacology—a cutting-edge computer-assisted drug development methodology—was unveiled. Investigating the underlying processes and complementary benefits of conventional herbal treatments is what it’s utilized for (!!! INVALID CITATION!!! [15]; Forthun et al., 2019). In this paper, a network pharmacology approach was integrated with molecular docking, simulation, GS-MS and FTIR that was utilized to investigate the active ingredients of *Papaver somniferum(Ganesh et al., 2023)*. It may expedite the drug discovery process by shedding new light on the molecular mechanisms behind Papaver somniferum’s anti-epileptic efficacy.

## 2. Materials and Methods

### 2.1 Screening of Chemical Constituents

All chemicals from Papaver somniferum were identified using literature and the KNApSAcK Core System database. All identified chemicals’ 2D and 3D structures were accessible through the PubChem database at https://pubchem.ncbi.nlm.nih.gov/. It is common practice to represent the molecular structure of compounds retrieved from the PubChem database using canonical SMILES. Papaver somniferum’s active compounds are evaluated pharmacokinetically using this succinct and consistent structure. Pharmacokinetics, which also includes the ADMET parameters, determines the likelihood of a drug molecule reaching the target protein in the body and its duration in the bloodstream. The characteristics of oral bioavailability (OB) and drug-likeness (DL) were investigated. Chemical characteristics such as solubility, permeability, and stability assess the likelihood of successfully developing a material into an oral medication, known as drug-likeness. A substance’s suitability for consumption depends on its body interaction. Oral bioavailability is the amount of epilepsy medication the body can use orally. Drug absorption and use are greatly affected by oral administration. Compounds that are drug-like (DL ≥ 0.18) and bioavailable orally (OB ≥ 30) were found to be good candidates for further research into how to treat epilepsy.

### 2.2 Screening for potential targets in Epilepsy disease

Initially, the canonical SMILES of the chosen probable constituents are extracted from PubChem, an extensive repository of chemical compounds. The SMILES strings are subsequently entered into SwissTargetPrediction, a technique utilized for forecasting the possible targets of small compounds. The tool utilizes SMILES strings to input molecular structure information and forecast probable interactions between the molecule and proteins or enzymes.Our analysis shows the drug’s epilepsy treatment mechanism. Human genes, products, and diseases are in GeneCards’ massive database. DisGeNET combines gene-disease data from multiple sources. By employing the keyword “epilepsy” in these databases, prospective targets associated with this ailment were chosen. The discovered targets from both databases were subsequently merged, presumably to generate a more exhaustive roster of candidate genes linked to the condition. Following the merging of the targets, any duplicated genes were eliminated. The targets’ standard names were acquired from UniProtKB. The Universal Protein Resource Knowledgebase, or UniProtKB, is an extensive database of information about protein sequences and their roles. By standardizing the names with UniProtKB, it is ensured that the targets can be easily and consistently identified. The organism for which the targets were selected and standardized is indicated as “Homo sapiens”. A Venn diagram is a visual representation that illustrates the correlation between different sets of genes related to “disorder” and “drug”. It is utilized to ascertain targets that are shared by both “disorder” and “drug.” The shared targets between the illness and the medicine should be regarded as prospective targets for further analysis of epilepsy.

### 2.3 Annotation of Gene Functions

The DAVID tool was utilized for conducting functional enrichment analysis on gene lists. Gene function annotation is determined using gene ontology analysis, which encompasses biological process (BP), cellular component (CC), and molecular function (MF). The KEGG database offers pathway analysis. The 123 venn genes underwent annotation and enrichment analysis using the DAVID program. The species chosen for this analysis was “Homo sapiens” (human). Significant pathways associated with epilepsy were discovered by analyzing enriched pathways with a probability value below 0.05. The Hiplot program was utilized to generate bubble plots of KEGG pathways and GO annotations, allowing for the visualization of enriched phrases linked to genes associated with epilepsy.

### 2.4. Network Pharmacology Analysis

#### 2.4.1. Construction of Compound-Target Networks

The active constituents of Papaver somniferum and the therapeutic target for epilepsy were utilized to construct the compound-target network in Cytoscape (version 3.8.2) (https://cytoscape.org/). Cytoscape is a costless instrument utilized for examining and illustrating biological networks composed of edges and nodes. Nodes in this context symbolize the chemical components and their specific genes and proteins associated with epilepsy, while edges symbolize the connection or interaction between them. The Cytoscape “network analyzer” application was utilized to ascertain the fundamental characteristics of the created network pertaining to epilepsy. The degree distribution was examined using a network analyzer to determine the number of connections that each node in the network possessed. The network was filtered based on the “degree” of each network node, which denotes the amount of connections that a certain node had in the epilepsy network.

#### 2.4.2 Prediction of the PPI Network and Hub Genes

The STRINGdatabase (version 11.0), developed by ELIXIR and located at the Welcome Genome Campus in Hinxton, Cambridgeshire, CB10 1SD, UK, was utilized to examine the connections among the therapeutic targets of epilepsy. The database can be accessed at https://string-db.org/. The organism selected was “*Homo sapiens*,” and a total score of 0.4 or greater was employed. The protein-protein interaction network associated with epilepsy was shown using Cytoscape. The CytoHubba plugin is utilized to identify the hub genes and nodes that have a higher degree inside the epilepsy network. The degree signifies that the targeted genes exhibit a substantial level of interconnectedness that is pertinent to epilepsy.

#### 2.4.3 Construction of Compound-Target-Pathway Networks

The network of target-compound-pathway associations linked to epilepsy was constructed using Cytoscape. This was achieved by utilizing the DAVID tool to evaluate the top 10 pathways, which were identified by KEGG pathway analysis of the targets and compounds associated with epilepsy.

### 2.5. GC-MS analysis

The phytochemical composition of Papaver somniferum was analyzed through the use of a Shimadzu QP2010 Plus GC-MS system. A thermal desorption system connected to the GC-MS was used to carry out the identification process. Seventy eV was the ionization voltage. A Restek column was used for the GC, which was run at 80–290°C. Both the injector and the interface had a temperature of 290°C. A flow rate of 1.2 ml/min of helium carrier gas was used to inject samples in split mode. Retention durations, mass spectra, and library matching to NIST and WILEY libraries were used to identify compounds. By comparing retention indices, mass spectra, and published data, active compounds were identified. Names, formulas, structures, and bioactivities of compounds could thus be determined.

### 2.6. FTIR

The extract’s unique functional groups were identified using the Fourier transform infrared (FTIR) technique. A molecule’s absorption spectrum can provide important structural details. A tiny quantity of dry potassium bromide (KBr) was mixed with Papaver somniferum extract. The mixture was meticulously blended in a mortar and compressed at a pressure of 6 bars for a duration of 2 minutes to create a thin disc made of KBr. Subsequently, the disk was inserted into a sample cup of a diffuse reflectance accessory. The IR spectrum was acquired using the Bruker Vertex 70 infrared spectrometer from Germany. The material underwent scanning spectroscopy from 4000 to 400 cm-1. The maximum values of the UV-VIS and FTIR spectra were recorded.

### 2.7 Molecular Docking

Potential active ingredients from Papaver somniferum that can target targets linked to epilepsy were found using molecular docking. Docking is used to predict how two molecules will align themselves in the best possible way when they are joined. It provides a more thorough comprehension of the molecular relationships between therapeutic protein targets and pharmaceutical compounds. The goal of the docking process was to confirm that the target predictions from the network pharmacology analysis were accurate. Compounds with higher docking scores were deemed to have better binding affinities and thus a greater likelihood of interacting with the therapeutic targets associated with epilepsy. The in silico molecular docking study facilitated the identification of potential drugs from Papaver somniferum that can strongly bind to crucial protein targets associated with epileptic seizures and related processes. Utilizing natural multi-target treatments provides a more secure option compared to existing anti-epileptic drugs, which may result in unwanted side effects. The integration of network pharmacology and molecular docking offers a comprehensive understanding of the molecular-level mechanisms through which drugs from Papaver somniferum may exert their potential anti-epileptic effects.

### 2.8 Molecular Dynamic Simulations

The validation of docking was conducted by employing molecular dynamics (MD) simulation with Desmond v3.6. The simulation employed the TIP3P model and an orthorhombic boundary box. The stability of protein-ligand interactions related to therapeutic targets for epilepsy was evaluated using the OPLS-2005 force field, with the addition of sodium ions. Before conducting the simulation, the protein-ligand system underwent minimization using the steepest descent (SD) and LBFGS algorithms. MD simulations were conducted using the Desmond software for a duration of 100 nanoseconds to examine the complexes resulting from the docking of potential compounds from Papaver somniferum with protein targets associated with epilepsy. This aided in confirming the stability of protein-ligand interactions that were identified in the docking experiments.

## 3. Results

### 3.1 Active Ingredients Screening

After identifying, filtering, and eliminating duplicates, a thorough compilation of active chemicals from Papaver somniferum was created by conducting a literature review and analyzing two distinct databases. After doing ADMET analysis on these compounds, we discovered 5 constituents that have the potential to be effective in treating epilepsy. These constituents were selected based on their drug-likeness (DL ≥ 0.18) and oral bioavailability (OB > 30) criteria. These drugs were deemed appropriate candidates for investigating the mechanisms of anti-epileptic activity and its interactions with therapeutic targets in epilepsy. The distinctive nature of these compounds spurred a thorough examination of their chemical structures using PubChem in order to provide insights into the anticipated interactions with protein targets associated with epilepsy. Table 1 represented the compounds with their ADMET properties.

**Table 1.** Selected bioactive compounds with DL, OB, BBB, Log P value.

| Sr. No | Compounds | DL | OB | BBB | Log P value | PubChem ID | Rotatable Bonds |
| --- | --- | --- | --- | --- | --- | --- | --- |
| 1 | Salutaridinol | 1.05 | 0.7714 | Yes | 1.74 | 5460042 | 2 |
| 2 | 7-O-Acetylsalutaridinol | 1.15 | 0.7286 | Yes | 2.2 | 5460163 | 4 |
| 3 | Scoulerine | 1.1 | 0.8143 | Yes | 2.41 | 22955 | 2 |
| 4 | Coclaurine | 0.89 | 0.6143 | Yes | 2.35 | 160487 | 3 |
| 5 | (-)-Codeinone | 0.73 | 0.5143 | Yes | 2.05 | 5459910 | 1 |
|  | (S)-Scoulerine | 1.1 | 0.8143 | Yes | 2.41 | 439654 | 2 |

### 3.2 Analysis of Network Pharmacology

### 3.3 Identifying Potential Targets

A total of 344 target genes of 5 active compounds were screened out through the SwissTargetPrediction database. Similarly, 5195 genes of epilepsy were screened from GeneCards and 4436 from DisGNet databases. 22 common targets of anti-epileptic treatment were selected that were potential targets of *Papaver somniferum*. These 22 shared targets between the compounds and epilepsy were considered for further analysis regarding their role in epileptic seizures and related pathways **as shown in figure 1**.

**Figure 1.**
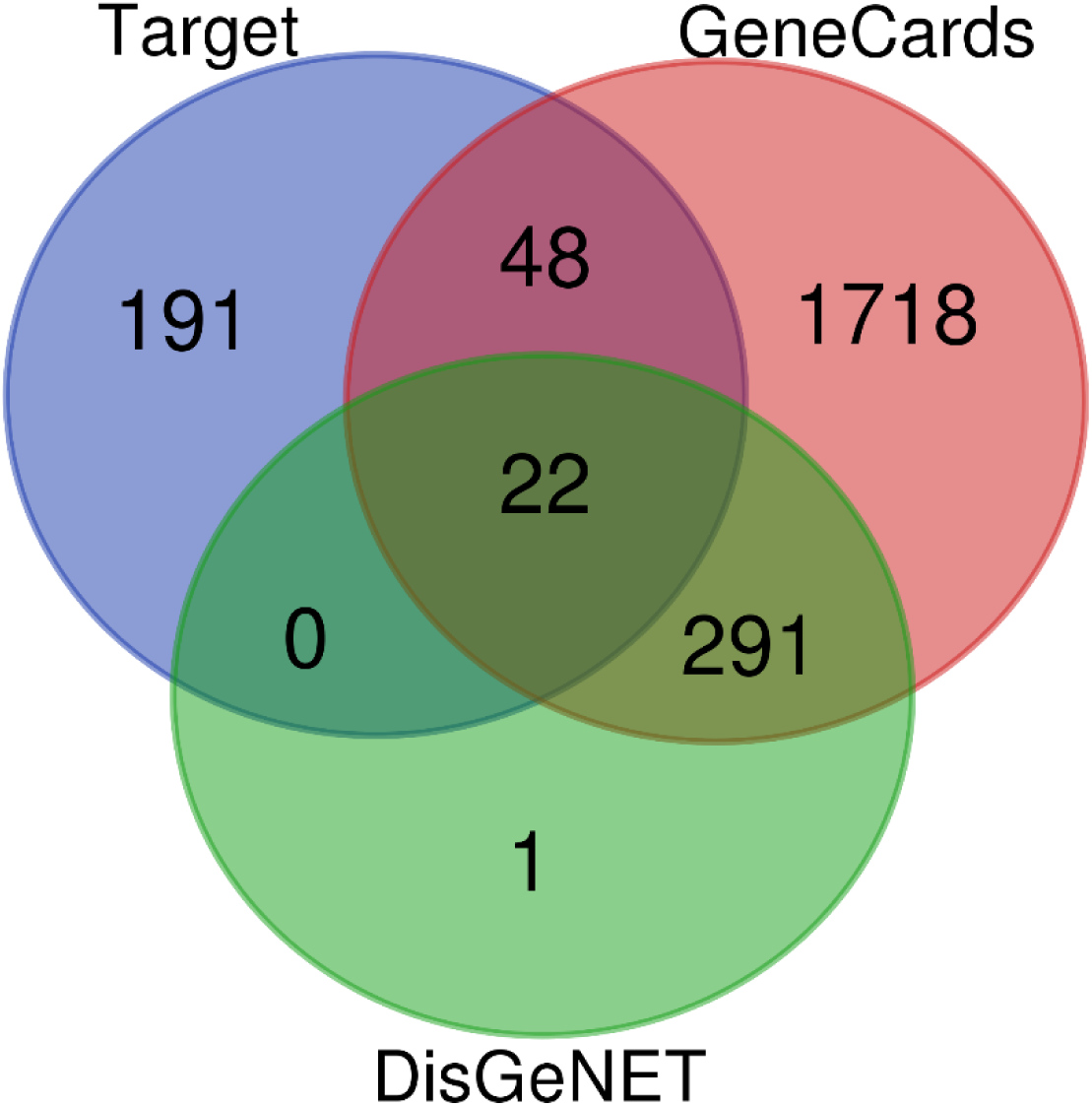
Venn diagram showing the common targets.

### 3.4 Compound target network

A network was constructed using Cytoscape to investigate the connection between active compounds derived from *Papaver somniferum* and possible targets associated with epilepsy, enabling the examination of their interaction processes. A network comprising 22 target genes linked to the treatment of epilepsy and 5 active constituents of *Papaver somniferum* shown in **table 2** was created. The use of the “network analyzer” tool showed that the compound-target network is made up of 1223 connections and 121 individual elements **as shown in figure 2**. The network analysis revealed the various ways in which the drugs affect epileptic seizures and the linked pathways at the molecular level.

**Figure 2.**
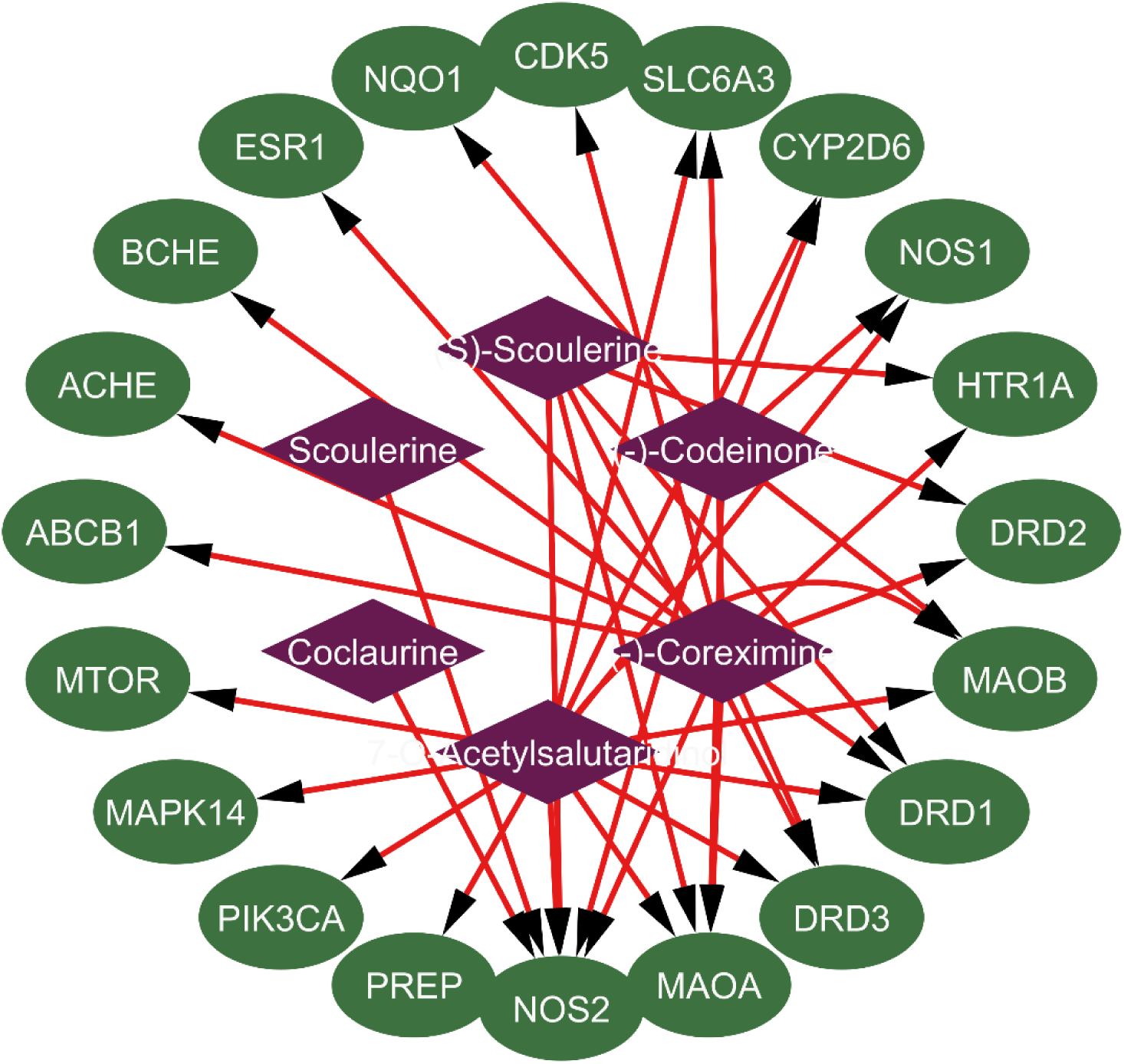
Compound-target network (dark color indicates active compounds; green colors indicate probable targets) connecting common targets of disease and chemicals.

The network consists of centrally positioned nodes colored in green, representing the components of Papaver somniferum. The surrounding nodes, colored in purple, represent the probable targets related with epilepsy. Moreover, the boundaries inside the network visually depict the connections between the compounds and their corresponding targets. The concentration of the 15 active components was analyzed in the compound-target network for epilepsy. The table displays alkaloids with varying degrees of concentration. Each of the 15 targets was selected for further molecular docking analysis. Based on the findings of the Target-compound network study, it has been seen that a solitary active molecule has the ability to engage with many targets linked to epilepsy. Additionally, it has been noted that the same target can simultaneously interact with multiple active components. This observation emphasizes the diverse and combined impacts of Papaver somniferum on the potential therapy of epilepsy by interacting with numerous key molecular targets.

### 3.5 PPI Network Analysis

Protein-protein interactions (PPI) are highly significant because to their wide range, adaptability, and accuracy. Examining PPI interactions reveals the links between the targets related to epilepsy as shown in **(Figure 5)**. This investigation offers significant insights into the interaction among many proteins implicated in epileptic seizures and associated pathways, illuminating the complex interactions within the biological system. Upon examining the PPI network in Cytoscape, it was noted that there were 22 nodes connected by 1223 edges associated with epilepsy targets and drugs **(Figure 3)**. The identification of central genes was conducted using the CytoHubba plugin. The network analysis toolkit comprises twelve topological techniques. The degree technique was employed to forecast hub genes within the epilepsy network. Greater degrees of connection indicate increased interaction of the targets with other elements within the context of epilepsy. This suggests that a higher degree may indicate a potential target that is implicated in epileptic pathways. The network has blue nodes that represent the top 10 hub genes, which are identified by their high degree values. The hub genes identified in this study include AKT1, IL6, HSP90AA, EGFR, ESR1, GSKB3, PTGS2, TLR4, APP, and ERBB2 **(Figure 4)**.

**Figure. 3.**
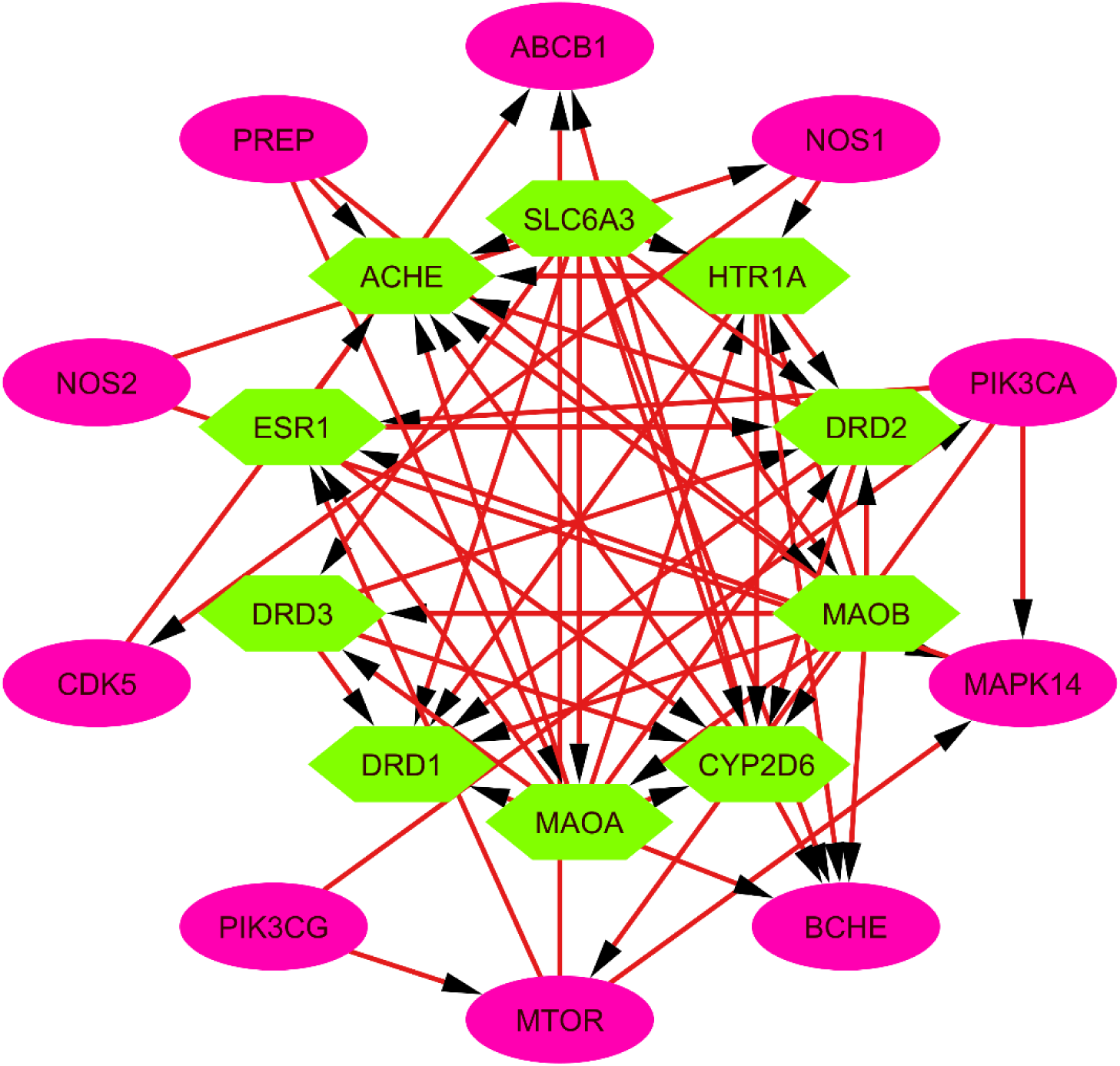
Analysis of 22 targets’ PPI networks. Ten targets are key nodes based on topological analysis (green nodes indicate additional possible targets, and central 22 nodes are hub genes with greater degrees).

**Figure. 4.**
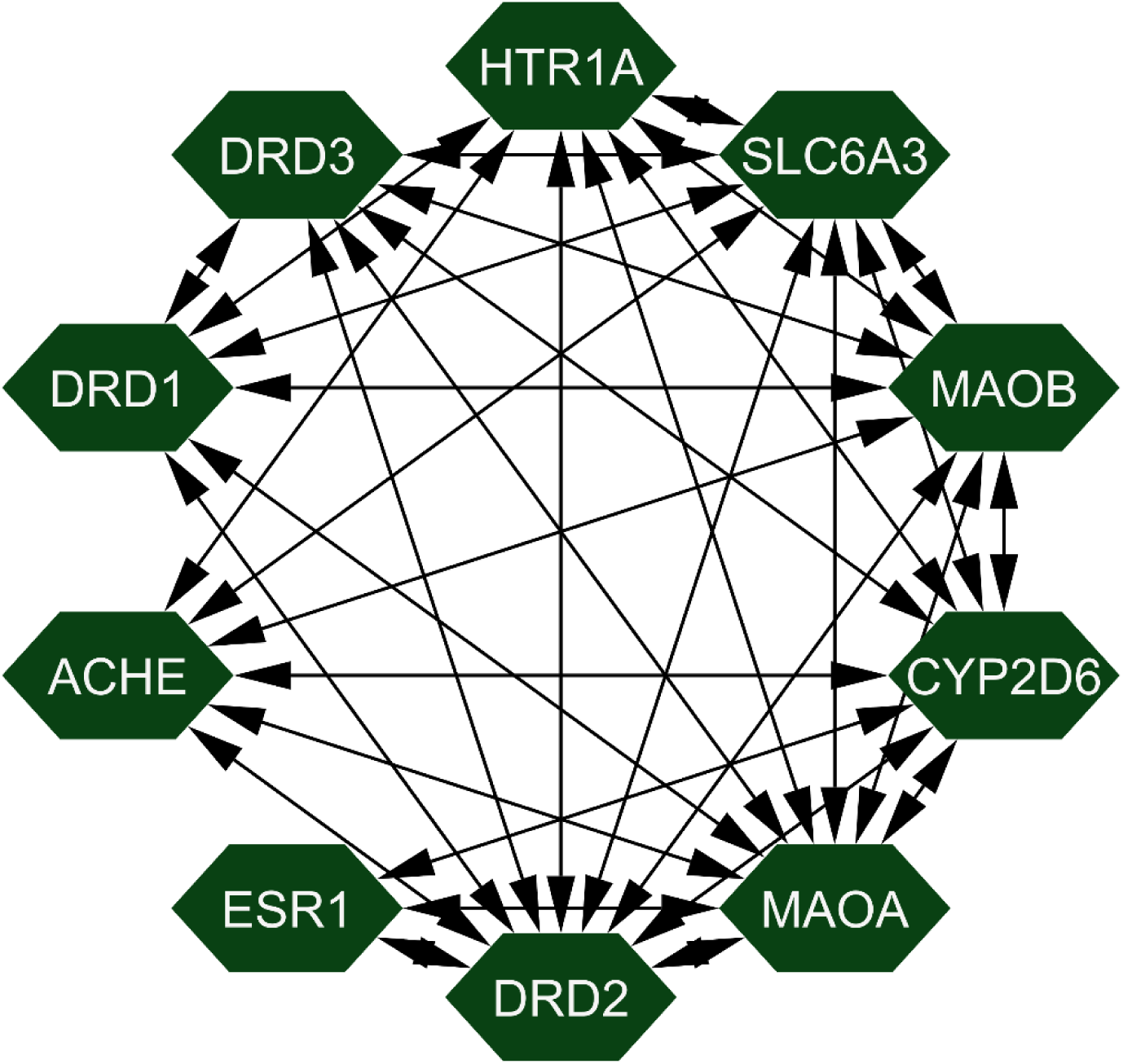
Top 10 hub genes responsible for controlling the pathways.

**Figure 5.**
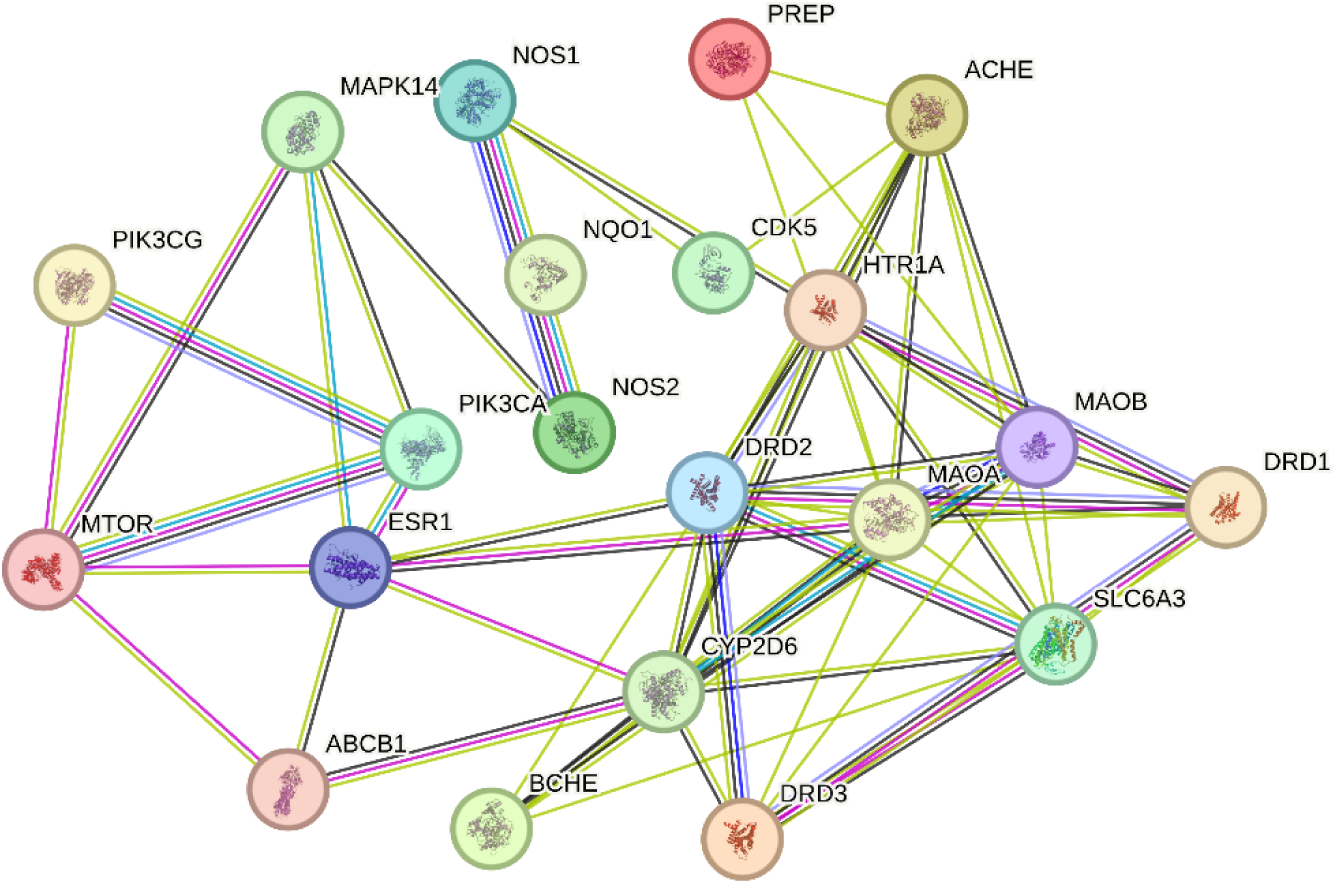
PPI network of 22 interconnected targets of epilepsy disease and *Papaver somniferum*.

### 3.7 KEGG Pathway and GO Analysis

An analysis of KEGG pathways and GO annotations was performed on the 123 possible targets related to the treatment of epilepsy. The GO analysis revealed a total of 518 biological processes (BP), 83 cellular components (CC), and 128 molecular functions (MF). The KEGG analysis found 141 pathways linked to epilepsy. The 20 best Gene Ontology (GO) annotations for Biological Process (BP), Cellular Component (CC), and Molecular Function (MF), along with the top 20 KEGG pathways, were selected based on a p-value threshold below 0.05. It was demonstrated that the annotations and pathways, which had a notable enrichment, were part of the targets linked to epilepsy. The Hiplot database was used to construct a bubble map that visually displayed the Gene Ontology and KEGG pathways. This study provided valuable insights into the primary functions and signaling mechanisms that are affected by the chemicals in their treatment of epilepsy, particularly at the systemic level.

### 3.8 Compound-Target-Pathway Network Analysis

Network analysis was utilized to investigate the putative anti-epileptic mechanism of Papaver somniferum. The DAVID analysis was used to identify the top 20 pathways that are enriched in epilepsy. Using Cytoscape, a network was then created to show the relationship between the targets, pathways, and compounds associated with epilepsy. The network consisted of 41 nodes and 85 edges, including eight active components derived from *Papaver somniferum*, 123 targets related to epilepsy, and 20 pathways **as shown in figure 6 and 7.** The degree values of each node were utilized to tailor its size and color. The targets demonstrated synchronization across many pathways implicated in epilepsy, establishing linked associations. The compounds found in Papaver somniferum may have a substantial impact on the possible anti-epileptic effects. These compounds possess the key characteristics of traditional Chinese medicine, such as targeting several aspects, containing multiple components, and acting through multiple pathways to combat a medical condition. This contributes to a comprehensive understanding of the systemic mechanisms by which Papaver somniferum can be used to treat epilepsy.

**Figure 6.**
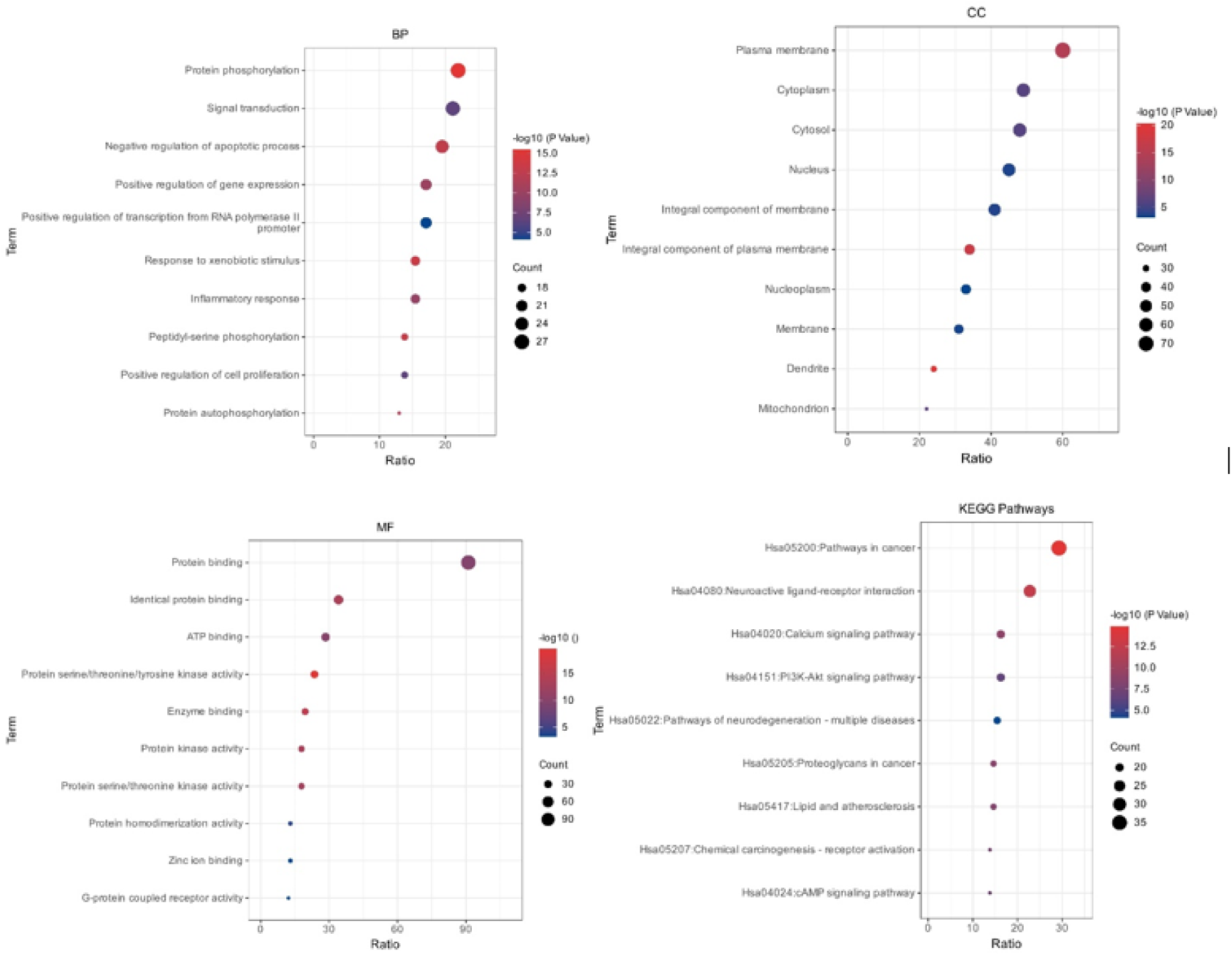
GO and KEGG pathway analysis bubble map (A) GO analysis’s biological process; (B) GO analysis’s cellular components; and (C) GO analysis’s molecular function.

**Figure. 7.**
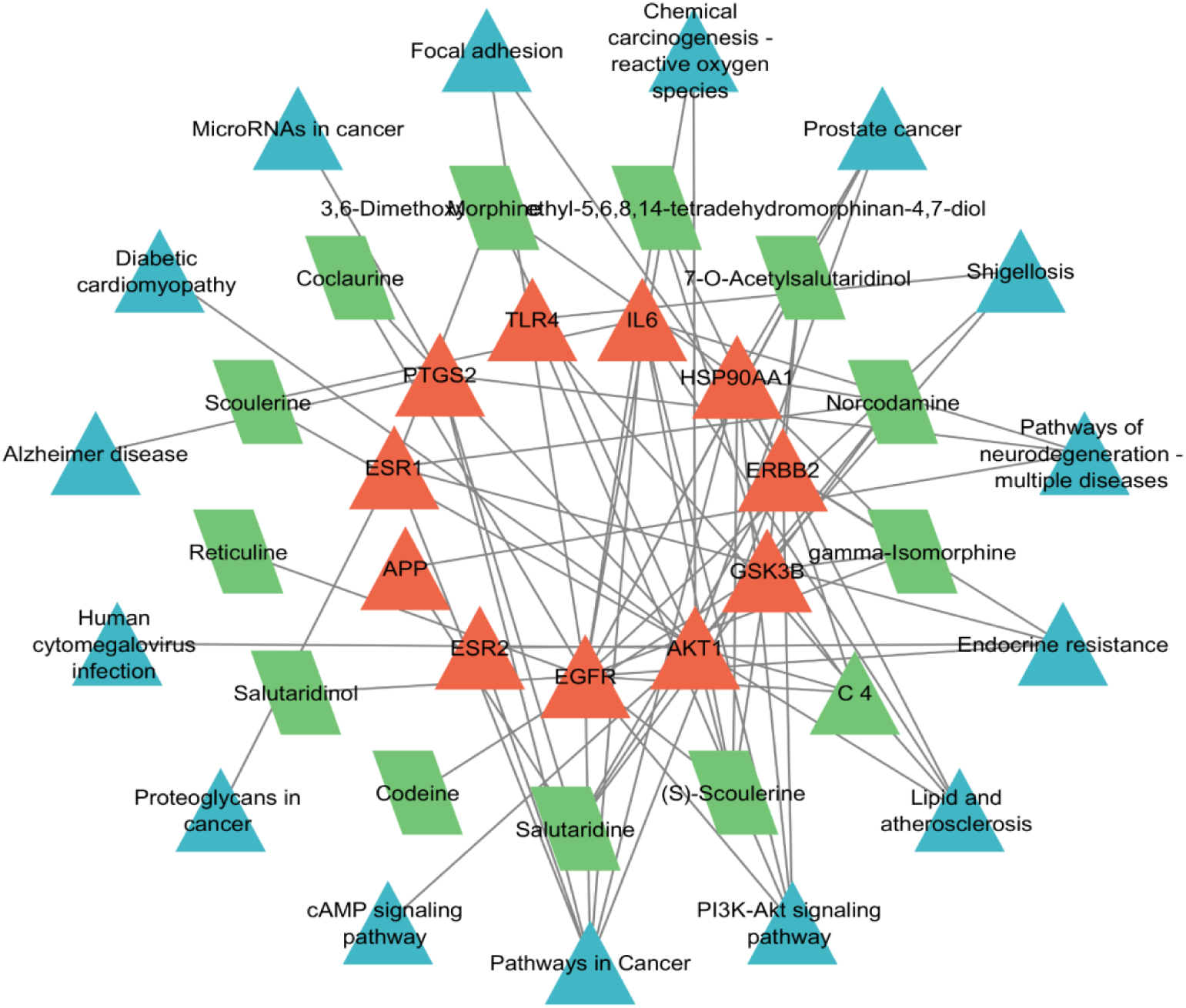
The network shows the relationship between active chemicals, target genes, and enrichment pathways. The size and color of the nodes indicate the level of interaction between them.

#### GC-MS Identification of plant extract phytochemicals

GC-MS study of Papaver somniferum methanolic plant extract revealed several chemicals. Table 3 shows 25 components in the methanolic extract. Papaver somniferum extract contains several alkaloids and related chemicals when extracted in methanol. Morphine and Codeine, with retention periods of 6.577 and 6.631 minutes, make up 4.98% and 9.39% of the extract. These compounds, known for their analgesic properties, are followed by Thebaine and Papaverine, present at 3.71% and 14.47% with retention times of 7.764 and 8.299 minutes, respectively. Papaverine, the most abundant alkaloid, is significant due to its vasodilatory effects. Noscapine and Narceine, with retention times of 9.358 and 9.808 minutes, are present at 1.91% and 8.97%, contributing to the extract’s profile with their anti-tussive and analgesic properties. Other notable compounds include Apomorphine (5.37% at 11.150 minutes) and Laudanosine (4.52% at 12.306 minutes), both of which have implications in neurological research. The presence of Protopine, Chelidonine, and Glaucine further diversifies the chemical profile, contributing to the extract’s potential therapeutic applications. Additionally, compounds like Berberine and Stylopine (15.114 and 15.889 minutes, 3.48% and 3.44%) highlight the extract’s role in traditional medicine. The analysis also identifies rare compounds such as Papaveraldine and Scoulerine (16.456 and 16.622 minutes, 1.26% and 3.52%), which are important for their unique bioactivity. Overall, this comprehensive profile underscores the complex pharmacological potential of Papaver somniferum leaves, reflecting both its historical medicinal use and its relevance in modern therapeutic contexts.

**Table 3:** Chemical Composition of Papaver somniferum Leaves (Methanolic Extract).

| Peak # | Retention Time | Area % | Molecular Formula | Name | IUPAC Name |
| --- | --- | --- | --- | --- | --- |

|  | (min) |  |  |  |  |
| --- | --- | --- | --- | --- | --- |
| 1 | 6.577 | 4.98 | C17H19NO3 | Morphine | (5α,6α)-7,8-didehydro-4,5-epoxy-17-methylmorphinan-3,6-diol |
| 2 | 6.631 | 9.39 | C18H21NO3 | Codeine | (5α,6α)-7,8-didehydro-4,5-epoxy-3-methoxy-17-methylmorphinan-6-ol |
| 3 | 7.764 | 3.71 | C19H21NO4 | Thebaine | (5α,6β)-7,8-didehydro-4,5-epoxy-3,6-dimethoxy-17-methylmorphinan |
| 4 | 8.299 | 14.47 | C20H21NO4 | Papaverine | (1S,3R)-6,7-dimethoxy-1-[(3S,4S)-3,4-dimethoxyphenyl]-3,4-dihydroisoquinoline |
| 5 | 9.358 | 1.91 | C22H23NO7 | Noscapine | 5-(4-ethenyl-1,2-dimethoxy-3-methylphenoxy)-4-methoxy-2,3-dihydroisoquinolin-2-one |
| 6 | 9.808 | 8.97 | C23H27NO4 | Narceine | 3,6-dimethoxy-5-[2-(7-methoxy-4-methyl-1,3-benzodioxol-5-yl)ethyl]-2H-1,3-benzodioxole-4-methanol |
| 7 | 11.150 | 5.37 | C17H21NO4 | Apomorphine | (R)-6-methyl-7,8-dihydro-6H-dibenzo[de,g]quinolin-7-ol |
| 8 | 12.306 | 4.52 | C20H21NO4 | Laudanosine | (S)-1-(3,4- |
|  |  |  |  |  | dimethoxybenzyl)-6,7-dimethoxyisoquinoline |
| **9** | 13.332 | 3.84 | C20H19NO5 | Protopine | 6,7-dimethoxy-4-(2-methylprop-1-en-1-yl)-3H-2-benzofuran-1-one |
| **10** | 13.423 | 1.33 | C21H21NO5 | Chelidone | (1R)-8-methyl-2,3,9,10-tetramethoxy-6H-[1,3]dioxolo[4,5-i]isoquinolin-6-one |
| **11** | 14.268 | 3.37 | C20H21NO4 | Papaveroline | 5-methoxy-7-methyl-6,7-dihydro-5H-dibenzo[de,g]quinolin-6-one |
| **12** | 14.584 | 1.72 | C21H25NO5 | Glaucine | 1-(3,4-dimethoxyphenyl)-4-methyl-3,4-dihydroisoquinolin-6,7-diol |
| **13** | 14.643 | 1.67 | C20H21NO4 | Sanguinarine | 6,7-dimethoxy-1-(3-methylbut-2-enyl)isoquinoline |
| **14** | 15.114 | 3.48 | C20H23NO5 | Berberine | (R)-6-methoxy-2,3-dihydro-9H-beta-carbolin-1-ylmethanol |
| **15** | 15.889 | 3.44 | C21H23NO4 | Stylophine | (S)-1-(3,4-dimethoxybenzyl)-7,8-dimethoxyisoquinoline |
| **16** | 16.456 | 1.26 | C22H21NO5 | Papaveraldine | (S)-1-(3,4-dimethoxybenzyl)-8,9-dimethoxy-1,2,3,4-tetrahydroisoquinoline |
| **17** | 16.622 | 3.52 | C21H23NO5 | Scoulerine | (S)-1-(3,4-dimethoxybenzyl)-6,7- |
|  |  |  |  |  | dimethoxy-1,2,3,4-tetrahydroisoquinoline |
| **18** | 16.991 | 3.25 | C19H19NO4 | Dehydrocrebanine | 6,7-dimethoxy-1-(3,4-dimethoxyphenyl)-2-methylisoquinolin-5(1H)-one |
| **19** | 17.082 | 4.78 | C22H23NO5 | Cryptopine | (S)-1-(3,4-dimethoxyphenyl)-3,4-dimethoxyisoquinoline |
| **20** | 17.312 | 3.27 | C21H25NO4 | Papaverubine | (R)-6-methoxy-1-methyl-2,3-dihydro-9H-beta-carbolin-9-one |
| **21** | 17.964 | 2.75 | C20H21NO4 | Canadine | 1-(3,4-dimethoxyphenyl)-6,7-dimethoxyisoquinolin-3-ol |
| **22** | 18.569 | 1.99 | C20H21NO4 | Dihydrochelerythrine | (R)-1-(3,4-dimethoxyphenyl)-1,2,3,4-tetrahydroisoquinoline |
| **23** | 19.136 | 1.41 | C21H23NO5 | Allocryptopine | 6,7-dimethoxy-1-(3,4-dimethoxyphenyl)-3,4-dihydroisoquinolin-5(1H)-one |
| **24** | 20.981 | 1.45 | C22H21NO5 | Emetine | (R)-1-(3,4-dimethoxyphenyl)-1,2,3,4-tetrahydroisoquinolin-6,7-diol |
| **25** | 21.414 | 4.14 | C20H19NO5 | Fumariline | (S)-1-(3,4-dimethoxyphenyl)-6,7-dimethoxy-1,2,3,4-tetrahydroisoquinoline |

The GC-MS chromatogram in **figure 8** of the methanol extract from Papaver somniferum plant exhibits a complex profile with multiple peaks, indicating the presence of various chemical constituents. The retention times range from approximately 6.0 to 23.0 minutes. The highest peak is observed at 8.202 minutes, signifying a compound with the highest abundance. Other significant peaks are noted at 6.832, 9.809, 11.152, 12.307, and 17.084 minutes, each representing different chemical compounds in the extract. The abundance of each peak varies, suggesting a diverse mixture of components within the sample. Peaks at retention times such as 13.332, 14.266, 15.111, and 16.624 minutes further indicate the presence of secondary metabolites. The baseline of the chromatogram shows minor fluctuations, suggesting some level of background noise or minor compounds in trace amounts. To identify and measure the different bioactive compounds present in the Papaver somniferum methanol extract, this chromatogram offers complete fingerprint. Understanding the phytochemical composition of the plant and its possible medicinal uses requires such thorough profiling. **Table 3** revealed the Chemical Composition of *Papaver somniferum* Leaves (Methanolic Extract).

**Figure 8.**
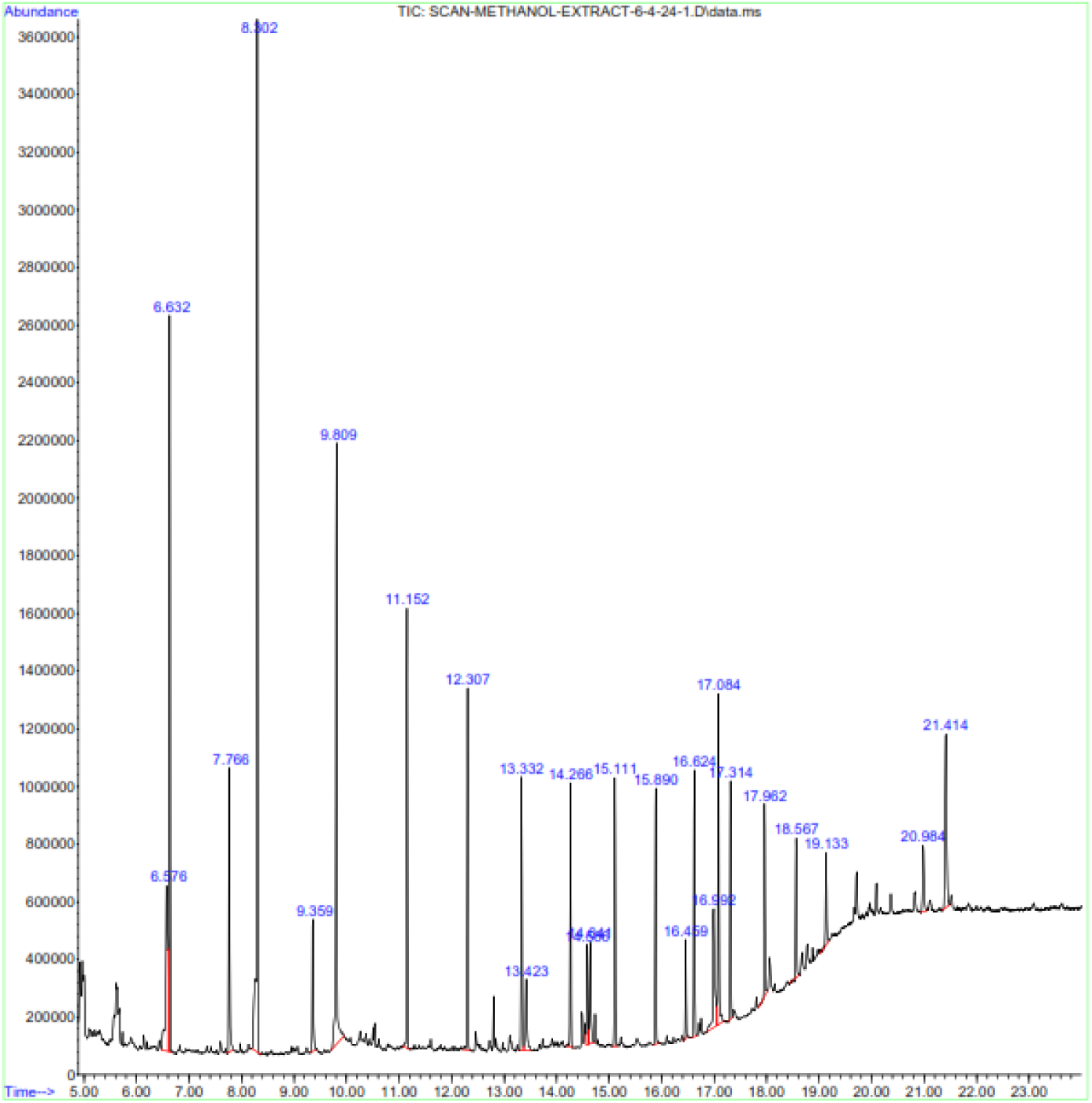
GC-MS chromatogram of methanol extract from Papaver somniferum plant. The chromatogram displays the total ion current (TIC) over the retention time. Major peaks are annotated with their respective retention times, representing various compounds present in the extract.

#### FTIR

The chemical bonds and functional groups in the dried leaf powder and leaf extract of Papaver somniferum were identified using FTIR spectroscopy. The analysis of the infrared absorption spectra allowed for the determination of the presence of specific bonds. The infrared absorption spectra were analyzed to determine the presence of specific bonds. A peak at 1026.34 cm−1 corresponds to the C-C skeletal vibration, indicating the presence of aromatic rings in the extract. The peak observed at 2848.28 cm−1 is attributed to the symmetric stretching of saturated (sp3) carbon, which is indicative of alkyl groups **as shown in figure 9**. The peak at 2916.41 cm−1 is associated with the asymmetrical stretching of CH3 groups, commonly found in alkaloids. Ester groups in phytoconstituents are indicated by the peak at 1894.84 cm−1, which represents the C=O stretching vibration of esters. Moreover, the peak at 2635.14 cm−1 indicates N-H stretching vibrations and suggests amine or amide groups. These absorption peaks indicate hydroxyl, carbonyl, aromatic, and other functional groups in Papaver somniferum extract phytoconstituents. **Table 4** represented the functional groups with their peak values using FTIR analysis.

**Figure 9:**
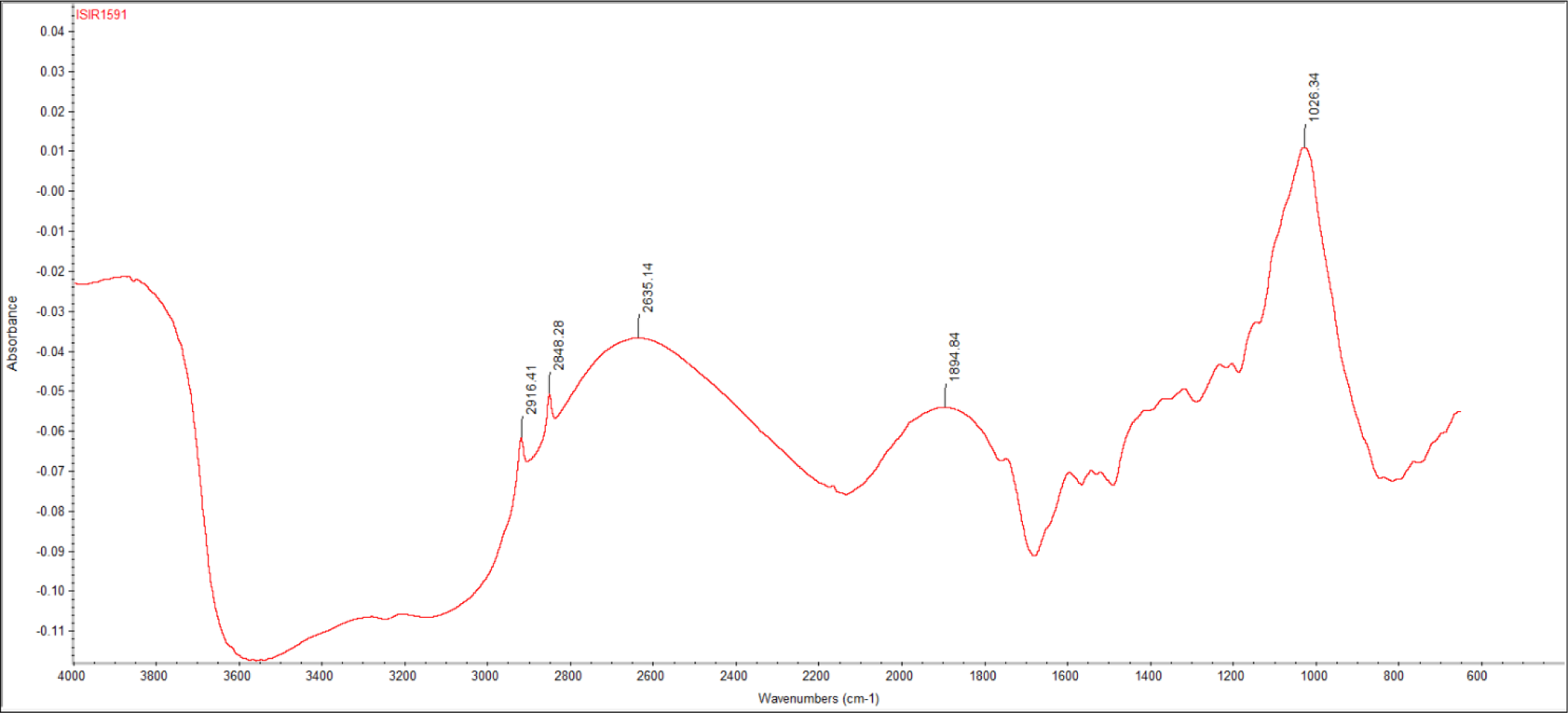
FTIR spectrum of dried leaf powder of Papaver somniferum.

**Table 4.** FTIR spectral peak values and functional groups of dried leaf powder of Papaver somniferum.

| Wave Numbers ($\text{cm}^{-1}$) | Functional Group | Predicted Compounds |
| --- | --- | --- |
| 2916.41 | CH<sub>3</sub> asymmetry stretching | Alkaloids |
| 2848.28 | C-H stretching vibration | Alkane |
| 2635.14 | N-H stretching vibration | Amine or Amide |
| 1894.84 | C=O of esters | Ester derivatives |
| 1026.34 | C-C skeletal vibration | Aromatic compounds |

### 3.9 Molecular Docking

The technique of molecular docking was employed to find prospective targets of components that have the ability to decrease the occurrence of epilepsy. The docking study successfully forecasted that the components would exhibit a robust affinity for the binding sites of the target proteins linked to epilepsy as shown in table 5. These genes were chosen by comparing the hub genes with the results obtained from the KEGG analysis of the epilepsy pathway. The molecular docking approach entailed the docking of 15 active components with 10 putative targets for epilepsy. In the field of molecular docking, a binding energy that is more negative indicates a stronger affinity between the ligand and receptor. The binding scores of all compounds and targets varied from −6 to −10 kcal/mole. Examining the interaction between these 10 targets associated with epilepsy is crucial for gaining a thorough understanding of the anti-epileptic effects of the active components **as shown in figure 10**. **Table 5** showed the docking table of the compounds with their targets and their binding energies.

**Figure 10.**
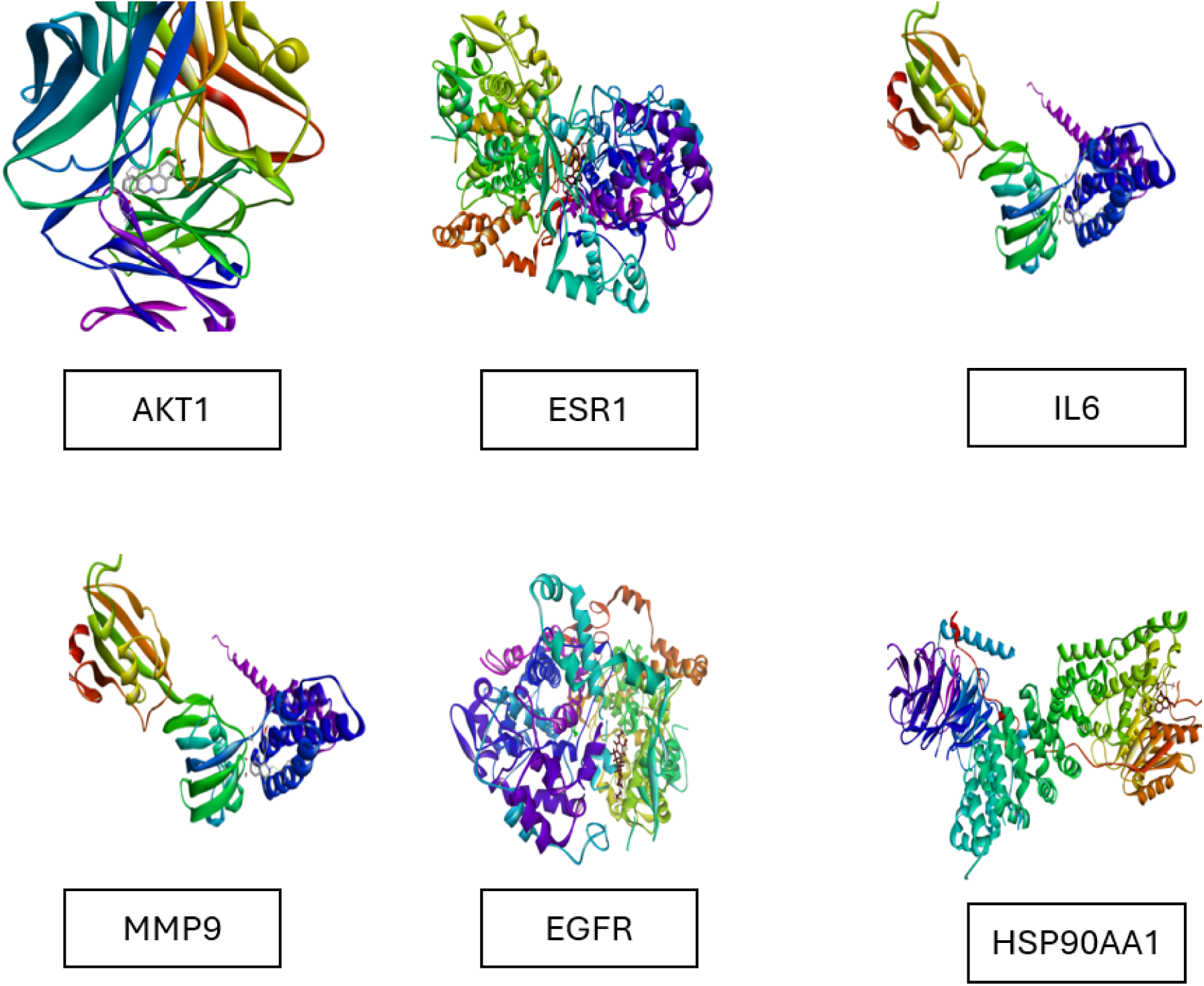
Docking complexes of *IL6, ESR1,MMP9, HSP90AA1*, and *AKT1* genes.

**Table 5.**
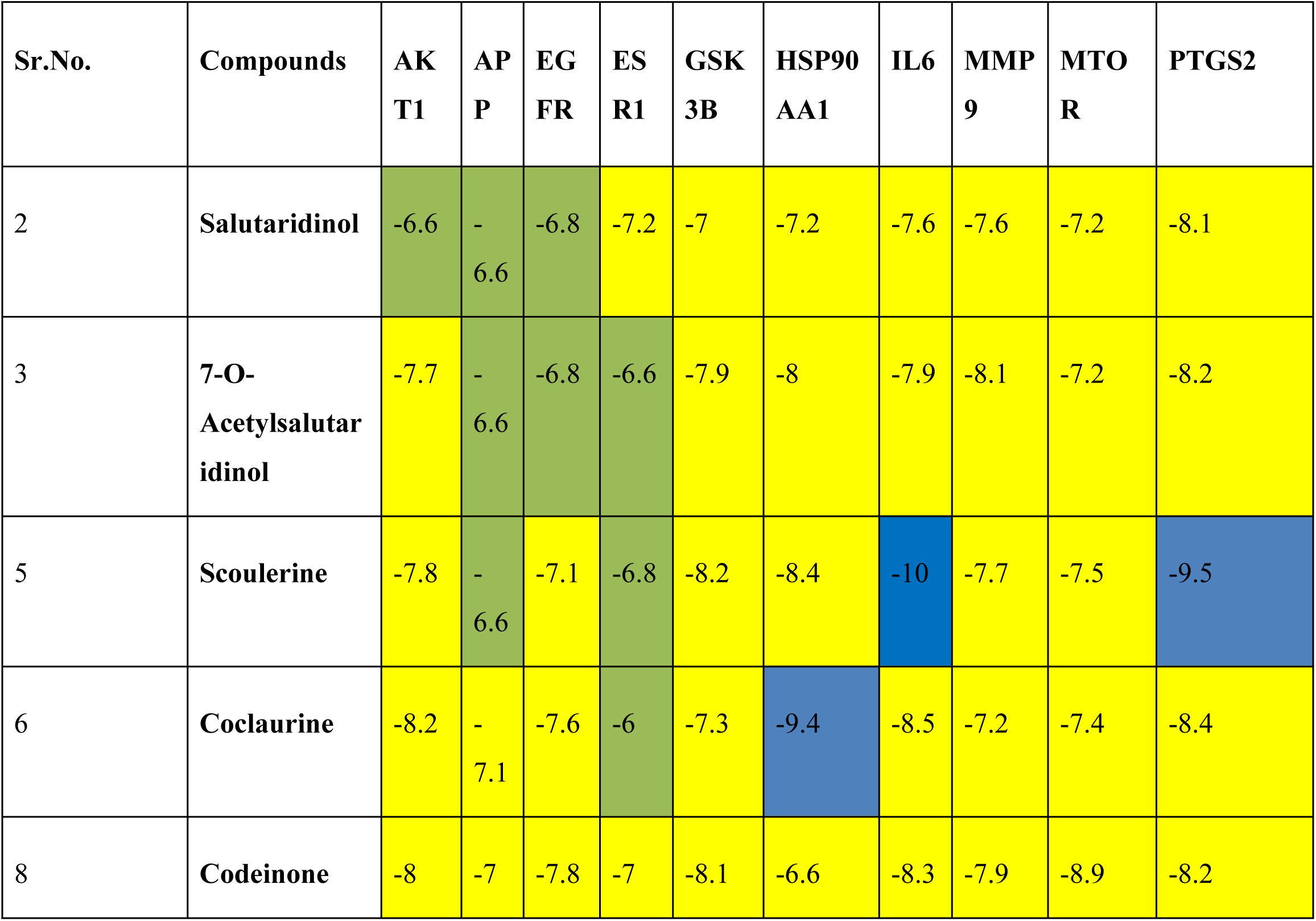
Molecular docking scores of active compounds with potent targets: Blue color; the highest docking score, yellow color; moderate docking score, green color; lower docking score.

| Sr.No. | Compounds | AK<br>T1 | AP<br>P | EG<br>FR | ES<br>R1 | GSK<br>3B | HSP90<br>AA1 | IL6 | MMP<br>9 | MTO<br>R | PTGS2 |
| --- | --- | --- | --- | --- | --- | --- | --- | --- | --- | --- | --- |
| 2 | Salutaridinol | -6.6 | -6.6 | -6.8 | -7.2 | -7 | -7.2 | -7.6 | -7.6 | -7.2 | -8.1 |
| 3 | 7-O-Acetylsalutaridinol | -7.7 | -6.6 | -6.8 | -6.6 | -7.9 | -8 | -7.9 | -8.1 | -7.2 | -8.2 |
| 5 | Scoulerine | -7.8 | -6.6 | -7.1 | -6.8 | -8.2 | -8.4 | -10 | -7.7 | -7.5 | -9.5 |
| 6 | Coclaurine | -8.2 | -7.1 | -7.6 | -6 | -7.3 | -9.4 | -8.5 | -7.2 | -7.4 | -8.4 |
| 8 | Codeinone | -8 | -7 | -7.8 | -7 | -8.1 | -6.6 | -8.3 | -7.9 | -8.9 | -8.2 |

### 3.10. Molecular Dynamics Simulation

The molecular dynamics (MD) simulation of the IL6 protein in complex with Scoulerine was conducted to understand the stability, flexibility, and binding interactions over a 100 ns period. The simulation provided detailed insights into the dynamic behavior of the protein-ligand complex, including structural deviations, residue fluctuations, and interaction profiles. These analyses are crucial for assessing the conformational stability and identifying key residues involved in binding, which can inform potential therapeutic implications and further experimental studies. The results presented here summarize the key findings from the RMSD, RMSF, and protein-ligand contact analyses conducted during the simulation.

#### 3.10.1. RMSD

Figure 11 shows a 100 ns molecular dynamics simulation of the IL6 protein with Scoulerine to evaluate the protein and ligand Root Mean Square Deviation (RMSD). In the initial 0-5 ns, the protein had a mean RMSD of 1.2 Å and a standard deviation of 0.3 Å, whereas the ligand had a mean RMSD of 3.5 Å and a standard deviation of 0.4. The protein’s mean RMSD grew to 1.5 Å (SD: 0.4 Å) and the ligand’s to 3.8 Å (SD: 0.5 Å) between 5 and 10 ns. From 10-15 ns, the protein and ligand mean RMSDs were 1.7 Å and 4.0 Å, respectively, with standard deviations of 0.5 and 0.6 Å. The protein and ligand mean RMSDs were 2.0 Å (SD: 0.6 Å) and 4.2 Å (SD: 0.6 Å) for the 15-20 ns timeframe. The protein had mean RMSDs of 2.3 Å (SD: 0.6 Å) while the ligand had 4.3 Å (SD: 0.7 Å) during the 20-25 ns timeframe. The mean RMSDs for the ligand and protein from 25-30 ns were 2.5 Å (SD: 0.7 Å) and 4.5 Å (SD: 0.8 Å), respectively. During the 30-35 ns interval, the protein and ligand mean RMSDs were 2.7 Å and 4.6 Å, respectively, with 0.8 Å and 0.8 Å standard deviation In the 35-40 ns range, protein and ligand mean RMSDs were 2.9 Å and 4.7 Å, respectively, with SDs of 0.8 Å and 0.9 Å. At 40-45 ns, the protein had a mean RMSD of 3.0 Å (SD: 0.9 Å) and the ligand had 4.8 Å (SD: 0.9 Å). Protein and ligand mean RMSDs within 45-50 ns were 3.2 Å (SD: 0.9 Å) and 4.9 Å (SD: 0.9 Å), respectively. The protein and ligand mean RMSDs in the 50-55 ns range were 3.3 Å and 5.0 Å, respectively, with SDs of 0.9 Å. The results were 3.4 Å (protein, SD: 1.0 Å) and 5.1 Å (ligand, SD: 1.0 Å) at 55-60 ns. The mean RMSDs for the protein and ligand during 60-65 ns were 3.5 Å and 5.2 Å, respectively, with an SD of 1.0 Å. Values were 3.6 Å (protein) and 5.3 Å (ligand) in the 65-70 ns range, with 1.0 Å SD for each. During the 70-75 ns timeframe, the protein and ligand had mean RMSDs of 3.7 Å and 5.4 Å, respectively, with 1.0 Å SDs Between 75 and 80 ns, the protein and ligand had mean RMSDs of 3.8 Å (SD: 1.1 Å) and 5.5 Å (SD: 1.0 Å). Between 80-85 ns, protein and ligand mean RMSDs were 3.9 Å and 5.6 Å, respectively, with SDs of 1.1 Å. In the 85-90 ns range, protein and ligand mean RMSDs were 4.0 Å and 5.7 Å, respectively, with SDs of 1.1 Å. In the 90-95 ns timeframe, the protein had a mean RMSD of 4.1 Å and the ligand 5.8 Å, with SDs of 1.1 Å. The mean RMSD of the ligand was 5.9 Å (SD: 1.1 Å) and the protein was 4.2 Å (SD: 1.1 Å) from 95-100 ns. Overall, the ligand had an average RMSD of 0.9 Å, while the protein had an average of 2.9 Å (SD: 0.8 Å). The ligand showed dynamic activity during the 100 ns simulation, indicating active binding interactions, but the IL6 protein remained structurally stable. Figure 11 exhibits RMSD. Figure 11 shows the RMSD.

**Figure. 11:**
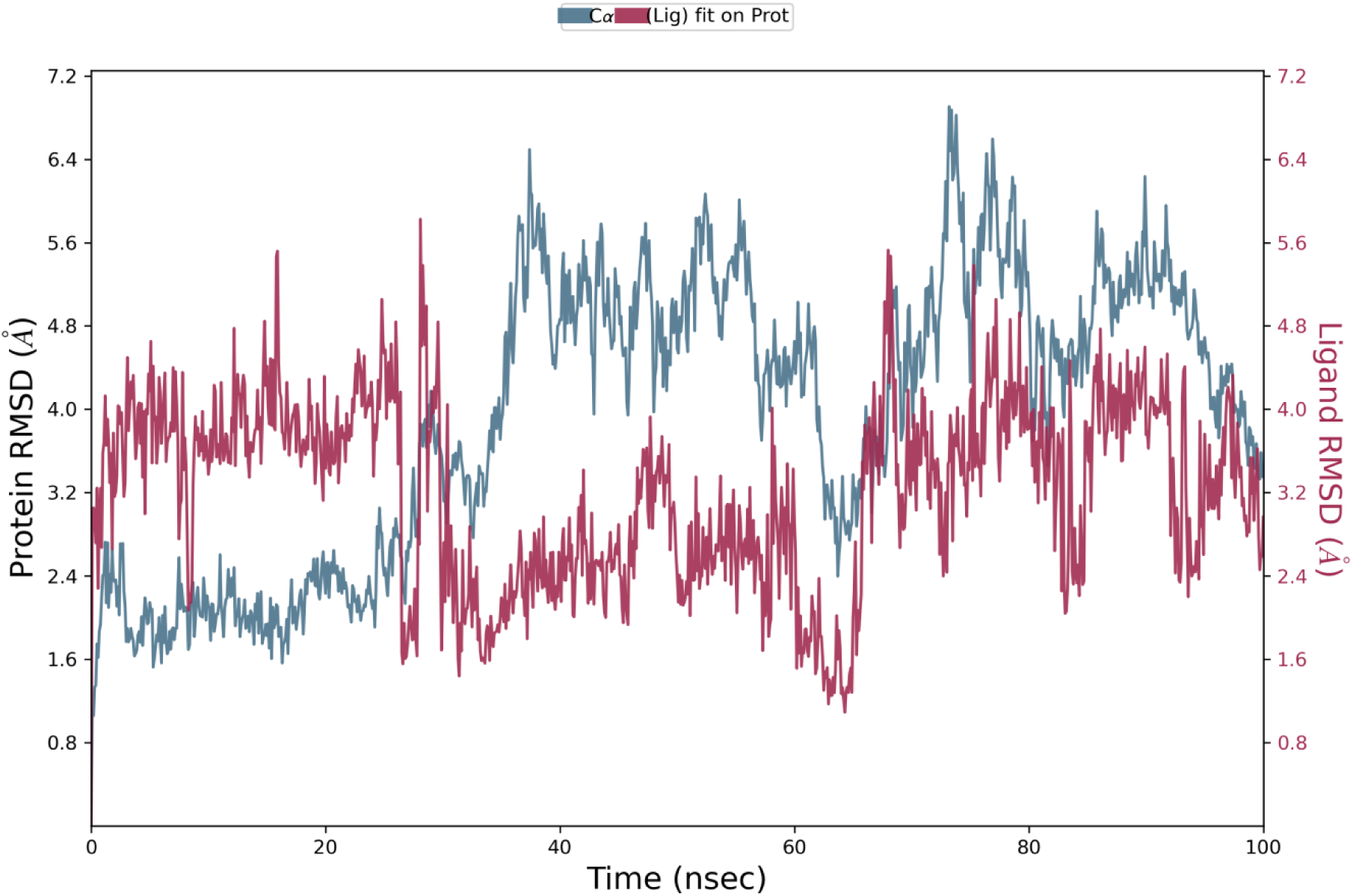
Protein and ligand RMSD over 100 ns MD simulation. The blue line represents the protein RMSD, and the red line represents the ligand RMSD. Data indicates protein stability and ligand fluctuations, demonstrating dynamic interactions within the IL6-Scoulerine complex.

#### 3.10.1. RMSF

The provided graph in figure 12 represents the Root Mean Square Fluctuation (RMSF) analysis of a protein-ligand complex, indicating the flexibility of residues within the protein structure. The degree of fluctuation that each residue experiences during the simulation is displayed by plotting the RMSF against the residue index. The y-axis displays the RMSF values in angstroms (é), and the x-axis shows the residue index. The RMSF values fluctuate around 1 to 2 Å for most residues, suggesting relatively stable regions with low flexibility. However, there are notable peaks, particularly towards the end of the residue index, indicating higher flexibility in these regions. The highest peak reaches approximately 9 Å, suggesting a significantly flexible or disordered region, possibly indicating areas of the protein that interact dynamically with the ligand or undergo conformational changes as shown in figure 13. Additionally, green vertical lines across the graph likely represent specific residues or points of interest, such as binding sites or regions of interaction with the ligand. These points coincide with varying RMSF values, suggesting differing degrees of flexibility and potential interaction dynamics.

**Figure 12:**
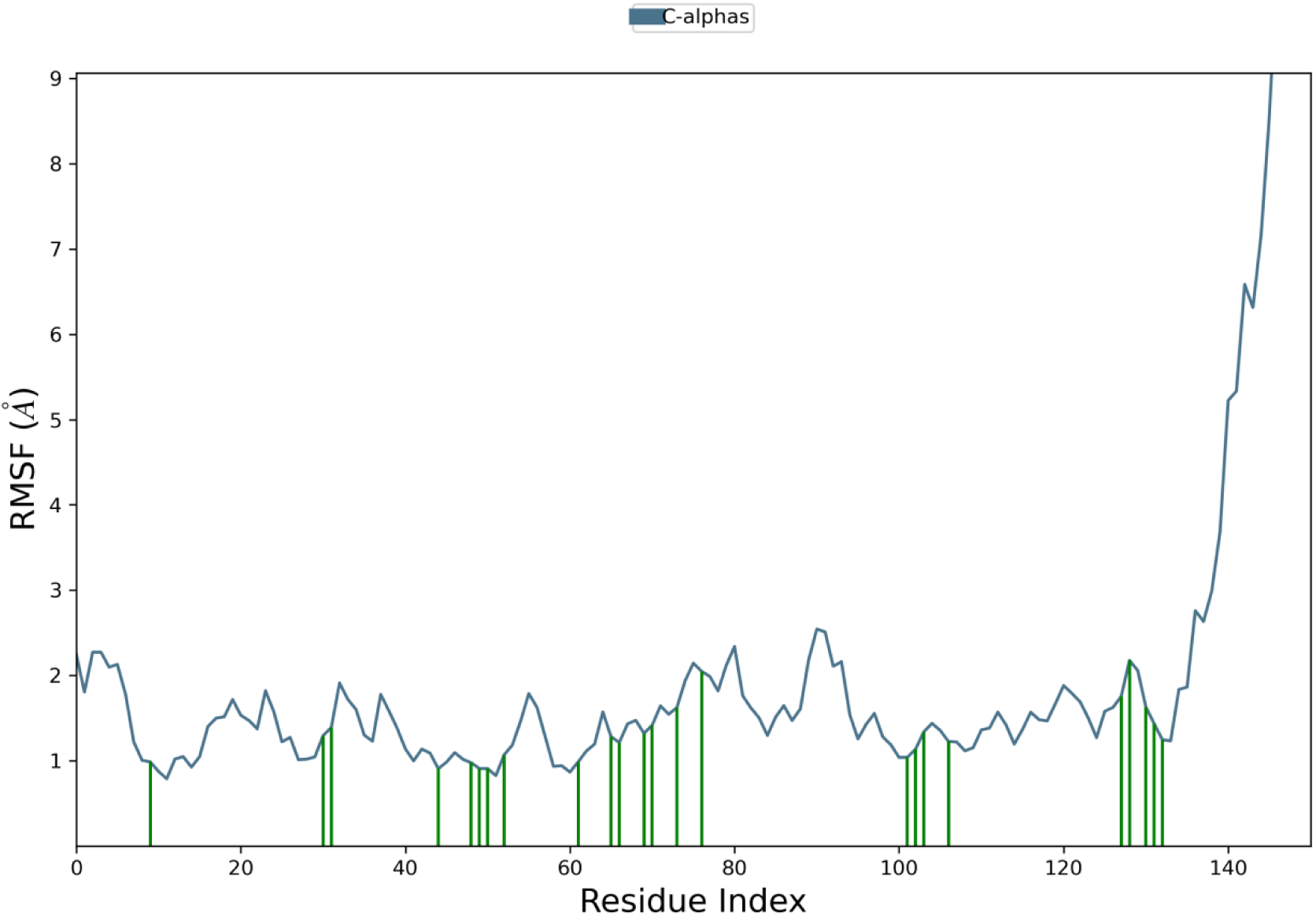
RMSF plot of the protein-ligand complex showing residue flexibility. The x-axis represents residue indices, and the y-axis shows RMSF values in angstroms (Å). Peaks indicate flexible regions, with the highest around 9 Å. Green lines mark specific residues of interest, highlighting interaction sites.

**Figure 13.**
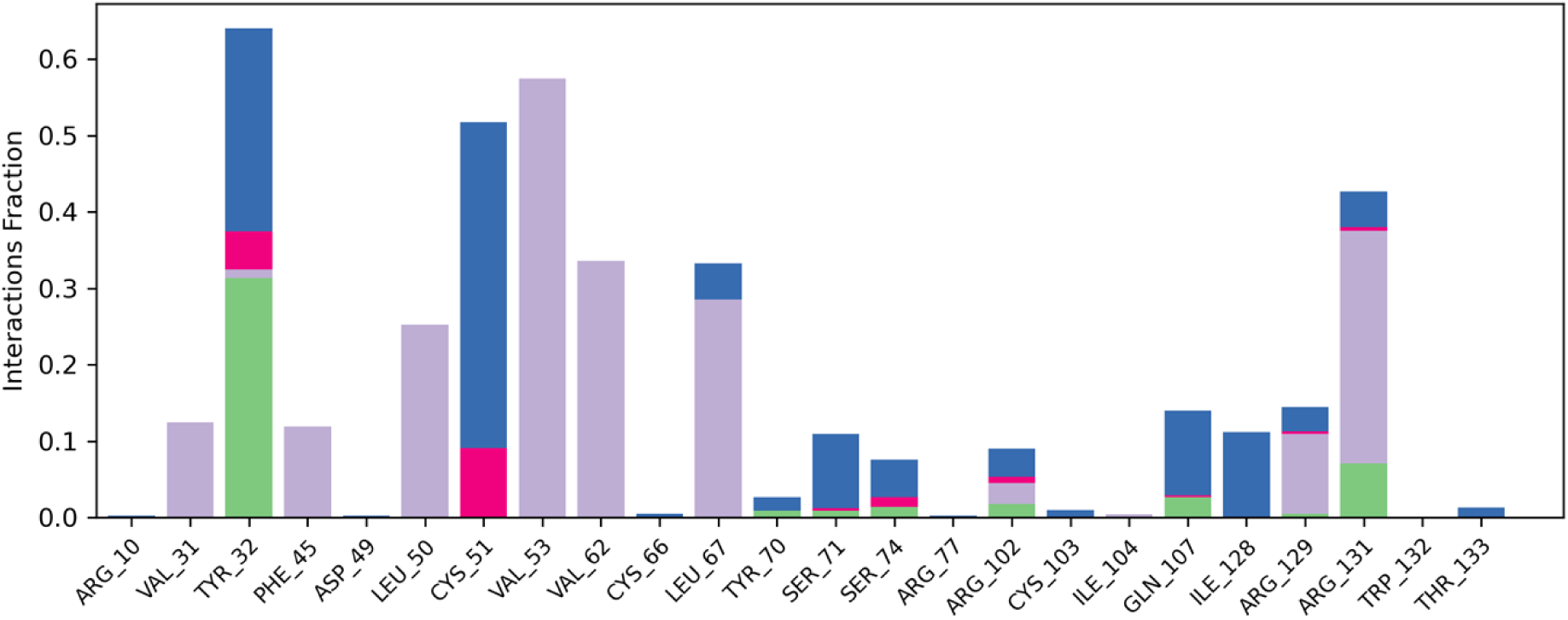
representing the protein ligand interaction map.

#### 3.10.2. Protein-Ligand Contacts

The provided histogram illustrates shown in figure 13 the interaction fractions of various amino acid residues in a protein-ligand complex, detailing the types of interactions these residues participate in. The y-axis shows the fraction of interactions for each amino acid residue, while the x-axis shows residues. Hydrophobic (purple), hydrogen bonds (green), ionic (red), and van der Waals forces (blue) are shown in each bar. Key observations include several residues exhibiting significant interaction fractions, indicating their critical roles in ligand stabilization and binding. For instance, TYR_32 exhibits substantial interaction fractions, predominantly involving hydrophobic interactions and hydrogen bonds, suggesting its crucial involvement in ligand stabilization. Similarly, CYS_51 shows a high interaction fraction with significant contributions from hydrophobic interactions and hydrogen bonds, underscoring its importance within the binding site. VAL_53 demonstrates notable interaction fractions with contributions from hydrophobic interactions and van der Waals forces, highlighting its role in maintaining the complex structure. ARG_131 displays significant interaction fractions, mainly involving hydrophobic interactions, hydrogen bonds, and ionic interactions, reflecting its multifaceted role in ligand binding. Other residues, such as LEU_50, VAL_62, and ARG_102, exhibit moderate interaction fractions, suggesting their supportive roles in stabilizing the protein-ligand complex through a mix of hydrophobic interactions and hydrogen bonds. Residues such as SER_71 and ARG_77 show lower interaction fractions, indicating they are less critical but still contribute to the overall stability of the complex through hydrophobic interactions and hydrogen bonds. Additionally, residues like PHE_45, ASP_49, and ILE_128 contribute to the interaction landscape with notable fractions, emphasizing the diverse nature of residue involvement in ligand binding. The histogram also highlights specific residues such as GLN_107, TRP_132, and THR_133, each contributing unique interaction profiles that include hydrophobic interactions and hydrogen bonds, further illustrating the complexity of the binding landscape. The presence of various interaction types within single residues, as depicted by the color variations, underscores the multifaceted nature of protein-ligand interactions. This comprehensive analysis of interaction fractions and types provides critical insights into the dynamic and intricate binding mechanisms within the protein-ligand complex. Understanding these detailed interaction patterns is crucial for elucidating the binding mechanisms and stability of the complex, which can inform drug design and protein engineering efforts. This detailed analysis provides valuable insights into the critical residues involved in ligand binding and their respective interaction contributions, essential for developing targeted therapeutic strategies and enhancing our understanding of protein-ligand dynamics.

#### Ligand Properties

The provided figure showcases various ligand properties over a 100-nanosecond (nsec) simulation, capturing dynamic changes and interactions of the ligand within the protein complex. The first panel illustrates the Root Mean Square Deviation (RMSD) of the ligand, with the y-axis representing the RMSD values in angstroms (Å). The consistent fluctuations between 0.30 and 0.45 Å indicate stable yet dynamic positioning of the ligand throughout the simulation. The second panel displays the Radius of Gyration (rGyr) of the ligand, indicating its compactness over time. The values oscillate around 0.85 Å, suggesting minor changes in the ligand’s compactness, which reflects its structural stability within the binding site.

The third panel represents the number of intramolecular hydrogen bonds (intraHB) within the ligand. Data oscillations between 0 and 2 show varied hydrogen bond creation and breaking, which may affect the ligand’s conformational stability and protein interaction. The ligand’s MolSA, ranging from 260 to 280 Å², is shown in the fourth panel. This reflects changes in the ligand’s accessible surface area, which may affect its protein binding site and solvent interactions. Fifth panel: solvent-accessible surface area (SASA), showing ligand solvent exposure. The SASA values vary between 60 and 90 Å², reflecting changes in the ligand’s orientation and interaction with the surrounding environment. The sixth panel shows the Polar Surface Area (PSA), which fluctuates between 175 and 180 Å². PSA is crucial for understanding the ligand’s solubility and permeability characteristics, with these fluctuations indicating variations in the polar regions exposed to the solvent. The histograms on the right side of each panel summarize the distribution of the respective properties over the simulation time. These distributions provide insights into the predominant states and fluctuations of the ligand properties throughout the simulation.

This comprehensive analysis of ligand properties, including RMSD, rGyr, intraHB, MolSA, SASA, and PSA, provides valuable insights into the ligand’s dynamic behavior and its interactions within the protein complex **as shown in** figure 14. Understanding these properties is crucial for elucidating the binding mechanisms, stability, and functional dynamics of the ligand, which can inform drug design and optimization efforts. This analysis highlights the importance of monitoring various ligand properties to gain a holistic understanding of its role and behavior in the protein-ligand complex.

**Figure. 14.**
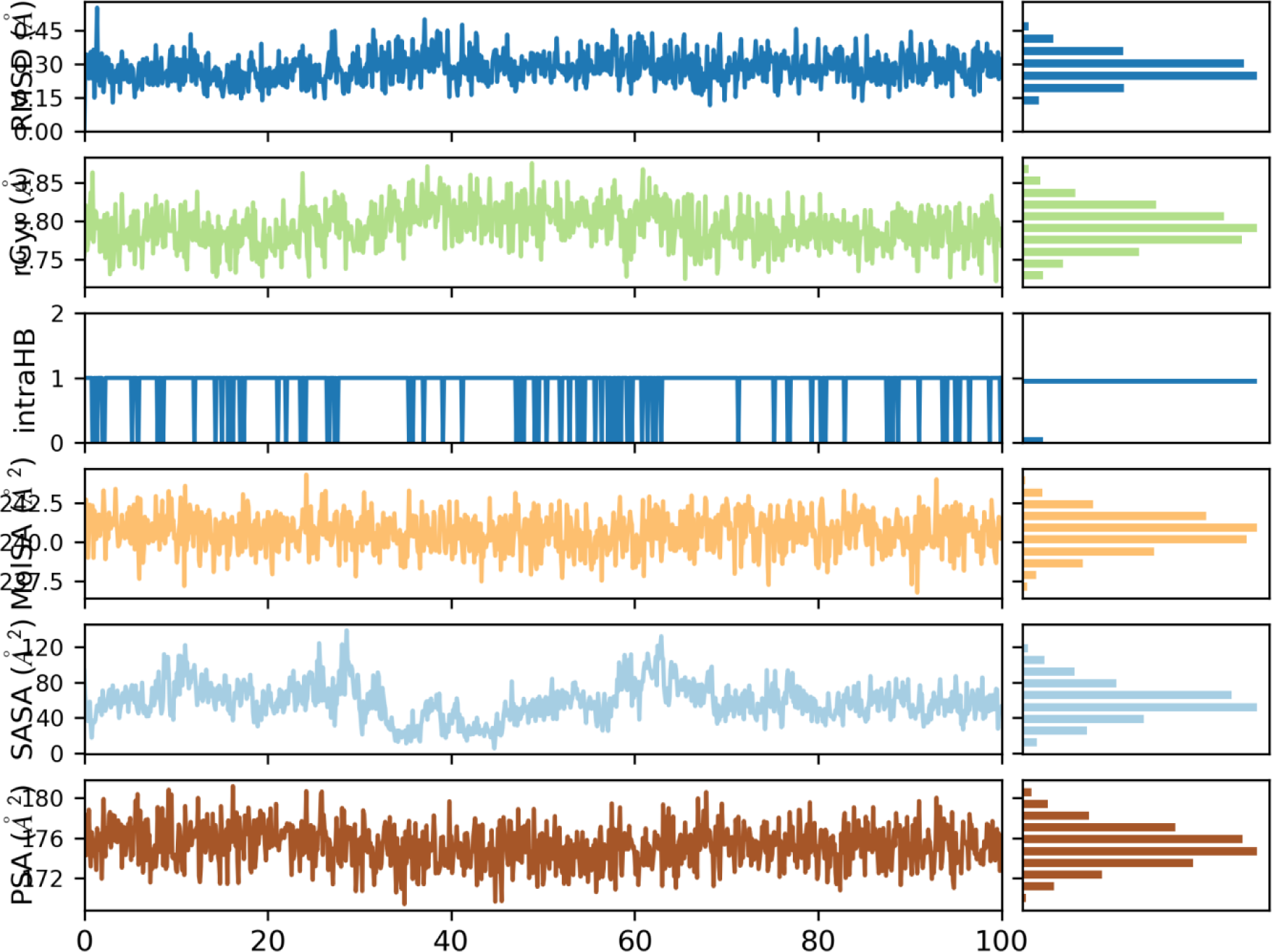
The provided figure various ligand properties over a 100-(nsec) simulation, capturing dynamic changes and interactions of the ligand within the protein complex.

## Discussion

The objective of this study was to explore, through a network pharmacology method, how Papaver somniferum may have anti-epileptic properties. Prior to commencing the research, we carried out an extensive review of databases and literature to detect and evaluate the active compounds in Papaver somniferum(Gasca-Perez et al., 2019; Ge et al., 2019). We performed an evaluation to determine the drug-likeness and oral bioavailability of the compounds in order to identify ones that are appropriate for managing epilepsy(Gomes et al., 2019). Studying the compositions of the substances and utilizing databases containing information on gene-disease links assisted in foreseeing potential protein targets associated with epilepsy(Gonzalez-Burgos et al., 2023).

A thorough literature review and database search identified and screened Papaver somniferum bioactive compounds. Drug-likeness and oral bioavailability were analyzed to find epilepsy-treating agents. Analysis of compound structures and gene-disease databases predicted epilepsy-related protein targets(He et al., 2021; Healy et al., 2023). Compounds and epilepsy were analyzed further based on shared targets. An analysis of gene ontology and pathway enrichment was conducted on the targets in order to obtain a deeper understanding of their activities and linkages. Interactions were visualized by constructing compound-target and protein-protein interaction networks. The process of molecular docking was employed to confirm the accuracy of target predictions(Huneif et al., 2022; Kanwal et al., 2022).

Five molecules - salutaridinol, 7-O-acetylsalutaridinol, scoulerine, coclaurine, and codeinone - were selected based on their drug-likeness and bioavailability. SwissTargetPrediction identified 344 targets for these drugs, whilst GeneCards and DisGeNet separately identified 5195 and 4436 genes related with epilepsy. 22 prevalent targets were chosen as potential mediators of Papaver somniferum’s anti-epileptic activities(Kodali et al., 2023; Kosem et al., 2021). The compound-target network unveiled a wide range of interactions between compounds and their effects on multiple targets. Through PPI network analysis, it was determined that there are 10 hub proteins that exhibit a high level of connectivity within the epilepsy network(Lamie et al., 2021; Maggi et al., 2022).

The gene ontology study revealed that the targets are implicated in more than 500 biological processes, 80 cellular components, and 120 molecular functions that are associated with epilepsy. The KEGG pathway analysis identified 141 pathways that were shown to be statistically significant, including pathways related to calcium signaling, MAPK signaling, and neuroactive ligand-receptor interactions(Mezzelani et al., 2023). The compound-target-pathway network illustrated the ability of Papaver somniferum to exert its effects across many paths. The molecular docking scores indicated a high affinity for binding between the drugs and hub targets. This confirms the accuracy of the network analysis forecasts and identifies possible mechanisms of anti-epileptic activity(Micheli et al., 2024; Navon et al., 2021).

This approach of integrative network pharmacology offers new insights into the complex mechanisms involving several targets that explain the traditional use of Papaver somniferum for treating epilepsy(Nogueira & Nunes, 2021; Pawlik et al., 2021). The identified bioactive substances, targets, and signaling pathways have the potential to serve as anti-epileptic components and interact at the molecular level. This enhances comprehension of the combined and interconnected impacts achieved through multi-component therapies such as herbal medications(Ramakrishnan et al., 2024; Rodrigues et al., 2023). The study confirms that Papaver somniferum is a highly promising source for the development of novel lead compounds with anti-epileptic properties. However, additional experimental verification is necessary to validate its therapeutic effectiveness and safety(Ruijs et al., 2022; Ruiz et al., 2020; Sabokrouh et al., 2023).

Ultimately, this study thoroughly examined the possible anti-epileptic mechanisms of Papaver somniferum by utilizing network pharmacology and molecular docking techniques. Virtual screening was used to identify five bioactive compounds that met drug-likeness requirements. Predictions were made regarding the shared targets of drugs and seizures, and the major proteins in the epilepsy network were identified(Samrani et al., 2023; Sharma et al., 2024; Sivakumar et al., 2022). The targets were analyzed using comprehensive gene ontology and pathway studies, which revealed valuable information about their biological activities and interactions. Network models provide a visual representation of the relationships among substances, targets, and pathways(Thokchom et al., 2024; Toniolo et al., 2020; Trombini et al., 2019). Molecular docking confirms target predictions by simulating the interactions between compounds and proteins(Wojcik et al., 2021). The study results demonstrate that the components of Papaver somniferum have several impacts on the pathways associated with epilepsy, and can be used as a basis for creating new treatments for epilepsy(Viereckl et al., 2022; Volmar et al., 2021; Walker et al., 2019). However, more experimental validations are required to convert the in silico findings into practical applications. In summary, the integrated network pharmacology technique is valuable for understanding the mechanisms of herbal medicines and expediting the process of drug discovery.

## Conclusion

The study used an integrated in silico framework to identify potential P. somniferum anti-epileptic leads. The literature and databases revealed twenty-five phytochemicals, including morphine, noscapine, and scoulerine. Bioinformatics analyses clarified their potential multi-targeted mechanisms of action against epilepsy. Network pharmacology outlined the systems-level interactions between the compounds and 124 potential targets. We predicted binding affinities between the compounds and ten important targets using molecular docking. During a 100-ns period, MD simulations confirmed that scoulerine and IL6 interacted dynamically and steadily. With its robust network centrality, stable binding conformations, and docking scores, scopularine became the front-runner. Its simulation sheds light on molecular recognition events by characterizing receptor flexibility, interactions between drug residues, and binding stability. The framework examined P. somniferum’s pleiotropic anti-seizure strategies at the atoms, proteins, networks, and systems levels. Important substances, such as scoulerine, require additional preclinical testing to support the rationale for anti-epileptic medication development. Focused experimental investigations using this multi-dimensional computational paradigm can guide the development of new anti-convulsants.

## Contribution

All authors contributed significantly to this study. Kiran Shahzadi led the study’s conceptualization, chemical analysis, and manuscript preparation. Danish Rasool conducted molecular docking and network pharmacology analyses. Faiza Irshad proofread and helps to perform GC-MS and FTIR for the project, also coordinated research activities, and critically revised the manuscript. Ammara Khalid contributed to molecular dynamics simulations and the analysis of RMSD and RMSF data. Sehar Aslam conceptualized and supervised the whole research work. Aisha Nazir assisted with GC-MS and FTIR data collection and analysis, and contributed to the manuscript writing. All authors reviewed and approved the final manuscript.

## Acknowledgment

Authors are thankful to University of Punjab for providing Server for MD simulation, GC-MS, and FTIR analyses

## References

Alves, C., Tamagno, W. A., Vanin, A. P., Pompermaier, A., & Barcellos, L. J. G. (2023). Cannabis sativa-based oils against aluminum-induced neurotoxicity. Sci Rep, 13(1), 9813. 10.1038/s41598-023-36966-9

Alyami, N. M., Abdi, S., Alyami, H. M., & Almeer, R. (2022). Proanthocyanidins alleviate pentylenetetrazole-induced epileptic seizures in mice via the antioxidant activity. Neurochem Res, 47(10), 3012–3023. 10.1007/s11064-022-03647-4

Andres-Mach, M., Szewczyk, A., Zagaja, M., Szala-Rycaj, J., Lemieszek, M. K., Maj, M., Abram, M., & Kaminski, K. (2021). Preclinical Assessment of a New Hybrid Compound C11 Efficacy on Neurogenesis and Cognitive Functions after Pilocarpine Induced Status Epilepticus in Mice. Int J Mol Sci, 22(6). 10.3390/ijms22063240

Araujo, G. F., Soares, L. O. S., Junior, S. F. S., Barreto de Carvalho, L. V., Rocha, R. C. C., Saint’Pierre, T., Hauser-Davis, R. A., Correia, F. V., & Saggioro, E. M. (2022). Oxidative stress and metal homeostasis alterations in Danio rerio (zebrafish) under single and combined carbamazepine, acetamiprid and cadmium exposures. Aquat Toxicol, 245, 106122. 10.1016/j.aquatox.2022.106122

Atmaca, E., Das, Y. K., Yavuz, O., & Aksoy, A. (2019). An evaluation of the levels of organochlorine compounds (OCPs and PCBs) in cultured freshwater and wild sea fish eggs as an exposure biomarker for environmental contamination. Environ Sci Pollut Res Int, 26(7), 7005–7012. 10.1007/s11356-019-04207-0

Baeeri, M., Rahimifard, M., Daghighi, S. M., Khan, F., Salami, S. A., Moini-Nodeh, S., Haghi-Aminjan, H., Bayrami, Z., Rezaee, F., & Abdollahi, M. (2020). Cannabinoids as anti-ROS in aged pancreatic islet cells. Life Sci, 256, 117969. 10.1016/j.lfs.2020.117969

Chantarat, N., Pe, K. C. S., Suppipat, K., Vimolmangkang, S., & Tawinwung, S. (2024). Effects of Cannabidiol on the Functions of Chimeric Antigen Receptor T Cells in Hematologic Malignancies. Cannabis Cannabinoid Res, 9(3), 819–829. 10.1089/can.2023.0108

Correa Basurto, A. M., Tamay Cach, F., Jarillo Luna, R. A., Cabrera Perez, L. C., Correa Basurto, J., Garcia Dolores, F., & Mendieta Wejebe, J. E. (2023). Hepatotoxic Evaluation of N-(2-Hydroxyphenyl)-2-Propylpentanamide: A Novel Derivative of Valproic Acid for the Treatment of Cancer. Molecules, 28(17). 10.3390/molecules28176282

DeFreitas, L., Fonseca Pego, A. M., Kronstrand, R., Lendoiro, E., de Castro-Rios, A., & Concheiro, M. (2022). Fast and Sensitive Method for the Determination of 17 Designer Benzodiazepines in Hair by Liquid Chromatography-Tandem Mass Spectrometry. J Anal Toxicol, 46(8), 852–859. 10.1093/jat/bkac044

Dervis, E., Karatay, K. B., Durkan, K., & Kilcar, A. Y. (2024). Radiolabeling of Zonisamide for a Diagnostic Perspective. Curr Radiopharm, 17(1), 91–98. 10.2174/0118744710249156231002115024

Domingos, L. B., Muller, H. K., da Silva, N. R., Filiou, M. D., Nielsen, A. L., Guimaraes, F. S., Wegener, G., & Joca, S. (2024). Repeated cannabidiol treatment affects neuroplasticity and endocannabinoid signaling in the prefrontal cortex of the Flinders Sensitive Line (FSL) rat model of depression. Neuropharmacology, 248, 109870. 10.1016/j.neuropharm.2024.109870

Engel, J., Jr., & Pitkanen, A. (2020). Biomarkers for epileptogenesis and its treatment. Neuropharmacology, 167, 107735. 10.1016/j.neuropharm.2019.107735

Forthun, R. B., Hellesoy, M., Sulen, A., Kopperud, R. K., Sjoholt, G., Bruserud, O., McCormack, E., & Gjertsen, B. T. (2019). Modulation of phospho-proteins by interferon-alpha and valproic acid in acute myeloid leukemia. J Cancer Res Clin Oncol, 145(7), 1729–1749. 10.1007/s00432-019-02931-1

Ganesh, S., Cortes-Briones, J., Schnakenberg Martin, A. M., Skosnik, P. D., D’Souza, D. C., & Ranganathan, M. (2023). Delta-9-Tetrahydrocannabinol, Cannabidiol, and Acute Psychotomimetic States: A Balancing Act of the Principal Phyto-Cannabinoids on Human Brain and Behavior. Cannabis Cannabinoid Res, 8(5), 846–856. 10.1089/can.2021.0166

Gasca-Perez, E., Galar-Martinez, M., Garcia-Medina, S., Perez-Coyotl, I. A., Ruiz-Lara, K., Cano-Viveros, S., Perez-Pasten Borja, R., & Gomez-Olivan, L. M. (2019). Short-term exposure to carbamazepine causes oxidative stress on common carp (Cyprinus carpio). Environ Toxicol Pharmacol, 66, 96–103. 10.1016/j.etap.2018.12.017

Ge, L., Cui, Y., Liu, B., Yin, X., Pang, J., & Han, J. (2019). ERalpha and Wnt/beta-catenin signaling pathways are involved in angelicin-dependent promotion of osteogenesis. Mol Med Rep, 19(5), 3469–3476. 10.3892/mmr.2019.9999

Gomes, T. B., Fernandes Sales Junior, S., Saint’Pierre, T. D., Correia, F. V., Hauser-Davis, R. A., & Saggioro, E. M. (2019). Sublethal psychotropic pharmaceutical effects on the model organism Danio rerio: Oxidative stress and metal dishomeostasis. Ecotoxicol Environ Saf, 171, 781–789. 10.1016/j.ecoenv.2019.01.041

Gonzalez-Burgos, I., Bainier, M., Gross, S., Schoenenberger, P., Ochoa, J. A., Valencia, M., & Redondo, R. L. (2023). Glutamatergic and GABAergic Receptor Modulation Present Unique Electrophysiological Fingerprints in a Concentration-Dependent and Region-Specific Manner. eNeuro, 10(4). 10.1523/ENEURO.0406-22.2023

He, Y., Jia, D., Du, S., Zhu, R., Zhou, W., Pan, S., & Zhang, Y. (2021). Toxicity of gabapentin-lactam on the early developmental stage of zebrafish (Danio rerio). Environ Pollut, 287, 117649. 10.1016/j.envpol.2021.117649

Healy, P., Verrest, L., Felisi, M., Ceci, A., Della Pasqua, O., & Consortium, G. (2023). Dose rationale for gabapentin and tramadol in pediatric patients with chronic pain. Pharmacol Res Perspect, 11(5), e01138. 10.1002/prp2.1138

Huneif, M. A., Alshehri, D. B., Alshaibari, K. S., Dammaj, M. Z., Mahnashi, M. H., Majid, S. U., Javed, M. A., Ahmad, S., Rashid, U., & Sadiq, A. (2022). Design, synthesis and bioevaluation of new vanillin hybrid as multitarget inhibitor of alpha-glucosidase, alpha-amylase, PTP-1B and DPP4 for the treatment of type-II diabetes. Biomed Pharmacother, 150, 113038. 10.1016/j.biopha.2022.113038

Jiao, W., Mi, S., Sang, Y., Jin, Q., Chitrakar, B., Wang, X., & Wang, S. (2022). Integrated network pharmacology and cellular assay for the investigation of an anti-obesity effect of 6-shogaol. Food Chem, 374, 131755. 10.1016/j.foodchem.2021.131755

Jiashuo, W. U., Fangqing, Z., Zhuangzhuang, L. I., Weiyi, J., & Yue, S. (2022). Integration strategy of network pharmacology in Traditional Chinese Medicine: a narrative review. J Tradit Chin Med, 42(3), 479–486. 10.19852/j.cnki.jtcm.20220408.003

Kanwal, M., Sarwar, S., Nadeem, H., Ata, A., Shah, F. A., Malik, S., Maqsood, S., & Miana, G. A. (2022). New pyrazolone derivatives, synthesis, characterization, and neuroprotective effect against PTZ-induced neuroinflammation in mice. Iran J Basic Med Sci, 25(12), 1424–1432. 10.22038/IJBMS.2022.62869.13912

Kodali, M., Jankay, T., Shetty, A. K., & Reddy, D. S. (2023). Pathophysiological basis and promise of experimental therapies for Gulf War Illness, a chronic neuropsychiatric syndrome in veterans. Psychopharmacology (Berl), 240(4), 673–697. 10.1007/s00213-023-06319-5

Kosem, A., Yucel, C., Titiz, A. P., Sezer, S., Neselioglu, S., Erel, O., & Turhan, T. (2021). Evaluation of serum thiol-disulphide homeostasis parameters as oxidative stress markers in epilepsy patients. Acta Neurol Belg, 121(6), 1555–1559. 10.1007/s13760-020-01410-6

Lamie, P. F., El-Kalaawy, A. M., Abdel Latif, N. S., Rashed, L. A., & Philoppes, J. N. (2021). Pyrazolo[3,4-d]pyrimidine-based dual EGFR T790M/HER2 inhibitors: Design, synthesis, structure-activity relationship and biological activity as potential antitumor and anticonvulsant agents. Eur J Med Chem, 214, 113222. 10.1016/j.ejmech.2021.113222

Li, L. Z., Zhou, C., Wang, P., Ke, Q. H., Zhang, J., Lei, S. S., & Li, Z. Q. (2023). Molecular mechanism of the effect of Gegen Qinlian decoction on COVID-19 comorbid with diabetes mellitus based on network pharmacology and molecular docking: A review. Medicine (Baltimore), 102(44), e34683. 10.1097/md.0000000000034683

Li, S., Chen, J., Hu, Y., & Ye, M. (2023). Editorial: Network pharmacology and AI. J Ethnopharmacol, 307, 116260. 10.1016/j.jep.2023.116260

Maggi, F., Morelli, M. B., Tomassoni, D., Marinelli, O., Aguzzi, C., Zeppa, L., Nabissi, M., Santoni, G., & Amantini, C. (2022). The effects of cannabidiol via TRPV2 channel in chronic myeloid leukemia cells and its combination with imatinib. Cancer Sci, 113(4), 1235–1249. 10.1111/cas.15257

Mezzelani, M., Peruzza, L., d’Errico, G., Milan, M., Gorbi, S., & Regoli, F. (2023). Mixtures of environmental pharmaceuticals in marine organisms: Mechanistic evidence of carbamazepine and valsartan effects on Mytilus galloprovincialis. Sci Total Environ, 860, 160465. 10.1016/j.scitotenv.2022.160465

Micheli, L., Lucarini, E., Nobili, S., Bartolucci, G., Pallecchi, M., Toti, A., Ferrara, V., Ciampi, C., Ghelardini, C., & Di Cesare Mannelli, L. (2024). Ultramicronized N-palmitoylethanolamine Contributes to Morphine Efficacy Against Neuropathic Pain: Implication of Mast Cells and Glia. Curr Neuropharmacol, 22(1), 88–106. 10.2174/1570159X21666221128091453

Navon, G., Novak, L., & Shenkar, N. (2021). Proteomic changes in the solitary ascidian Herdmania momus following exposure to the anticonvulsant medication carbamazepine. Aquat Toxicol, 237, 105886. 10.1016/j.aquatox.2021.105886

Nogueira, A. F., & Nunes, B. (2021). Acute and chronic effects of diazepam on the polychaete Hediste diversicolor: Antioxidant, metabolic, pharmacologic, neurotoxic and behavioural mechanistic traits. Environ Toxicol Pharmacol, 82, 103538. 10.1016/j.etap.2020.103538

Pawlik, M. J., Miziak, B., Walczak, A., Konarzewska, A., Chroscinska-Krawczyk, M., Albrecht, J., & Czuczwar, S. J. (2021). Selected Molecular Targets for Antiepileptogenesis. Int J Mol Sci, 22(18). 10.3390/ijms22189737

Ramakrishnan, S., Singh, T., & Reddy, D. S. (2024). Protective Activity of Novel Hydrophilic Synthetic Neurosteroids on Organophosphate Status Epilepticus-induced Chronic Epileptic Seizures, Non-Convulsive Discharges, High-Frequency Oscillations, and Electrographic Ictal Biomarkers. J Pharmacol Exp Ther, 388(2), 386–398. 10.1124/jpet.123.001817

Rodrigues, P., Guimaraes, L., Carvalho, A. P., & Oliva-Teles, L. (2023). Carbamazepine, venlafaxine, tramadol, and their main metabolites: Toxicological effects on zebrafish embryos and larvae. J Hazard Mater, 448, 130909. 10.1016/j.jhazmat.2023.130909

Ruijs, T. Q., Koopmans, I. W., de Kam, M. L., van Esdonk, M. J., Koltzenburg, M., Groeneveld, G. J., & Heuberger, J. (2022). Effects of Mexiletine and Lacosamide on Nerve Excitability in Healthy Subjects: A Randomized, Double-Blind, Placebo-Controlled, Crossover Study. Clin Pharmacol Ther, 112(5), 1008–1019. 10.1002/cpt.2694

Ruiz, C. E., Manuguerra, S., Curcuraci, E., Santulli, A., & Messina, C. M. (2020). Carbamazepine, cadmium chloride and polybrominated diphenyl ether-47, synergistically modulate the expression of antioxidants and cell cycle biomarkers, in the marine fish cell line SAF-1. Mar Environ Res, 154, 104844. 10.1016/j.marenvres.2019.104844

Sabokrouh, A., Sadeghi Motlagh, B., & Atabi, F. (2023). Study of anticancer effects of platinum levetiracetam and levetiracetam via cancer biomarkers genes expression on HepG2 cell line. Mol Biol Rep, 50(11), 9431–9439. 10.1007/s11033-023-08890-8

Samrani, L. M. M., Dumont, F., Hallmark, N., Bars, R., Tinwell, H., Pallardy, M., & Piersma, A. H. (2023). Nervous system development related gene expression regulation in the zebrafish embryo after exposure to valproic acid and retinoic acid: A genome wide approach. Toxicol Lett, 384, 96–104. 10.1016/j.toxlet.2023.07.005

Sharma, P., Kumari, A., Singh, P., Srivas, S., Thakur, M. K., & Hemalatha, S. (2024). Pyrus pashia fruit extract and its major phytometabolite chrysin prevent hippocampal apoptosis and memory impairment in PTZ-kindled mice. Nutr Neurosci, 27(8), 836–848. 10.1080/1028415X.2023.2276575

Sivakumar, P. M., Prabhakar, P. K., Cetinel, S., R, N., & Prabhawathi, V. (2022). Molecular Insights on the Therapeutic Effect of Selected Flavonoids on Diabetic Neuropathy. Mini Rev Med Chem, 22(14), 1828–1846. 10.2174/1389557522666220309140855

Thokchom, S. K., Indracanti, N., Khanna, A., & Indraganti, P. K. (2024). Safety evaluation of 5-hydroxytryptophan and S-(2-aminoethyl)isothiouronium bromide hydrobromide on rodent lungs. Indian J Pharmacol, 56(1), 28–36. 10.4103/ijp.ijp_176_23

Toniolo, S., Sen, A., & Husain, M. (2020). Modulation of Brain Hyperexcitability: Potential New Therapeutic Approaches in Alzheimer’s Disease. Int J Mol Sci, 21(23). 10.3390/ijms21239318

Trombini, C., Hampel, M., & Blasco, J. (2019). Assessing the effect of human pharmaceuticals (carbamazepine, diclofenac and ibuprofen) on the marine clam Ruditapes philippinarum: An integrative and multibiomarker approach. Aquat Toxicol, 208, 146–156. 10.1016/j.aquatox.2019.01.004

Viereckl, M. J., Krutsinger, K., Apawu, A., Gu, J., Cardona, B., Barratt, D., & Han, Y. (2022). Cannabidiol and Cannabigerol Inhibit Cholangiocarcinoma Growth In Vitro via Divergent Cell Death Pathways. Biomolecules, 12(6). 10.3390/biom12060854

Volmar, M. N. M., Cheng, J., Alenezi, H., Richter, S., Haug, A., Hassan, Z., Goldberg, M., Li, Y., Hou, M., Herold-Mende, C., Maire, C. L., Lamszus, K., Fluh, C., Held-Feindt, J., Gargiulo, G., Topping, G. J., Schilling, F., Saur, D., Schneider, G., … Glass, R. (2021). Cannabidiol converts NF-kappaB into a tumor suppressor in glioblastoma with defined antioxidative properties. Neuro Oncol, 23(11), 1898–1910. 10.1093/neuonc/noab095

Walker, D. I., Perry-Walker, K., Finnell, R. H., Pennell, K. D., Tran, V., May, R. C., McElrath, T. F., Meador, K. J., Pennell, P. B., & Jones, D. P. (2019). Metabolome-wide association study of anti-epileptic drug treatment during pregnancy. Toxicol Appl Pharmacol, 363, 122–130. 10.1016/j.taap.2018.12.001

Wojcik, P., Gegotek, A., Zarkovic, N., & Skrzydlewska, E. (2021). Disease-Dependent Antiapoptotic Effects of Cannabidiol for Keratinocytes Observed upon UV Irradiation. Int J Mol Sci, 22(18). 10.3390/ijms22189956

